# Genetic control of pod dehiscence in domesticated common bean: Associations with range expansion and local aridity conditions

**DOI:** 10.1101/517516

**Authors:** Travis A. Parker, Jorge C. Berny Mier y Teran, Antonia Palkovic, Judy Jernstedt, Paul Gepts

**Affiliations:** University of California, Department of Plant Sciences, 1 Shields Avenue, Davis, CA 95616-8780

**Keywords:** dirigent, aridity tolerance, GWAS, local adaptation, pod shattering

## Abstract

**Significance:** Plant domestication has radically modified crop morphology and development. Nevertheless, many crops continue to display some atavistic characteristics that were advantageous to their wild ancestors, such as pod dehiscence (PD). Domesticated common bean (*Phaseolus vulgaris*), a nutritional staple for millions of people globally, shows considerable variation in PD. Here, we identified multiple genetic regions controlling PD in common bean grown throughout geographically distributed lineages. For example, on chromosome Pv03, *PvPdh1* shows a single base-pair substitution that is strongly associated with decreased PD and expansion of the crop into northern Mexico, where the arid conditions promote PD. The environmental dependency and genetic redundancy explain the maintenance of atavistic traits under domestication. Knowledge of PD genetics will assist in developing aridity-adapted varieties.

**Abstract:** A reduction in pod dehiscence (PD) is an important part of the domestication syndrome in legumes, including common bean. Despite this, many modern dry bean varieties continue to suffer yield reductions due to dehiscence, an atavistic trait, which is particularly problematic in hot, dry environments. To date, the genetic control of this important trait has been only partially resolved. Using QTL mapping and GWAS, we identified major PD QTLs in dry beans on chromosomes Pv03, Pv05, Pv08, and Pv09, three of which had not been described previously. We further determined that the QTL on chromosome Pv03, which is strongly associated with PD in Middle American beans, includes a dirigent-like candidate gene orthologous to *Pod dehiscence 1 (Pdh1)* of soybean. In this gene, we identified a substitution in a highly conserved amino acid that is unique to PD-resistant varieties. This allele is associated with the expansion of Middle American domesticated common beans into the arid environments of northern Mexico, resulting in a high allelic frequency in the domesticated ecogeographic race Durango. The polygenic redundancy and environmental dependency of PD resistance may explain the maintenance of this atavistic characteristic after domestication. Use of these alleles in breeding will reduce yield losses in arid growing conditions, which are predicted to become more widespread in coming decades.

## Introduction

Effective seed dispersal is vital for spermatophytes. In the Fabaceae, the third largest family of flowering plants (1), seed dispersal is typically mediated by the explosive dehiscence (“shattering”) of pods at maturity. This mechanism is effective for the propagation of plants in the wild, but is associated with reduced yield and harvest constraints for cultivated crops. As such, there has been selection against pod dehiscence (PD), which continues to this day. A reduction in PD has been a central part of the domestication syndrome of many domesticated pulses. Anatomical differences are associated with some, but not all variation in PD in these species (2, 3). Reviews of the developmental genetics related to PD are available (4, 5).

In soybean (*Glycine max* (L.) Merr.), a domestication-related reduction in PD is mediated by the *NAC* family transcription factor *SHAT1-5* (6). A further reduction in PD is controlled by *Pod dehiscence 1* (*Pdh1) (7). Pdh1* encodes a dirigent-like gene related to lignin synthesis. This mutation is associated with the expansion of soybeans into arid regions. *Pdh1* is highly expressed in obliquely oriented fibers lining the soybean pod walls and has a minimal effect on gross pod anatomy (8).

Common bean (*Phaseolus vulgaris* L.) is the foremost grain legume for direct human consumption and is a dietary staple for hundreds of millions of people worldwide (9). Common bean diverged into distinct Middle American and Andean gene pools approximately 87,000 years before present, well before the first human migrations into the Americas (10). Subsequently, human populations independently domesticated members of each gene pool, making up two of at least seven domestication events in the genus *Phaseolus* (11) and 41 domestication events in the Fabaceae (12). Each of the two major domesticated gene pools of common bean is divided into several ecogeographic races. In the Middle American domesticated gene pool, it is important to single out race Durango, which includes varieties from arid, higher altitude regions of northern Mexico, and race Mesoamerica, from the warmer, humid lowlands of southern Mexico and Central America (13).

*Phaseolus vulgaris* can be separated into two economic groups: snap beans, grown for pods as a vegetable, and dry beans, grown for grain. Dry beans produce fibrous pods, which can be easily separated from seeds during threshing. In snap beans, selections in the 19^th^ century led to “stringless” varieties with extreme PD resistance and very little pod suture fiber deposition (14). Stringless varieties now dominate the snap bean market, but stringlessness is absent in dry beans. Using a recombinant inbred (RI) population derived from stringless cv. ‘Midas’ and wild accession G12873, Koinange *et al.* (15) identified a major pod fiber QTL on linkage group Pv02 (16). This gene, called *Stringless (St),* maps near the common bean ortholog of *INDEHISCENT (PvIND*), but there is a lack of complete co-segregation between the loci and no causal polymorphism is known to exist in the *PvIND* sequence (17). Rau *et al.* (18) used QTL mapping to identify a segregating locus on Pv05 in the Midas x G12873 genetic background. Despite this, a comprehensive evaluation of the genetic basis of PD in diverse germplasm has not yet been conducted and no molecular polymorphisms with a potential causal effect on PD have been described.

In the research reported here, we used high-precision phenotyping techniques, both in an RI population and diversity panels, to identify PD QTLs in common bean. A locus underlying one of the major QTLs was sequenced to identify possible causal polymorphisms. We found orthologous mechanisms that regulate pod dehiscence in this species. We were further able to identify associations between PD and local environmental conditions. Alleles identified in this study will be valuable for developing common bean varieties suited to the increasingly arid climatic conditions of coming decades.

## Results

### Anatomical analysis of developing pods

Clear differences in pod anatomy were found between domesticated snap bean, domesticated dry bean, and wild common bean (Fig. 1). Wild beans produce a lignified wall fiber layer (LFL) in the pods that is thicker than the bundle cap layer, while the LFL is greatly reduced in domesticated varieties. Stringless snap beans have weakly lignified vascular bundle sheaths (VS) at the suture, with a reduction in the number of lignified cells and the extent of secondary cell wall deposition in each cell. In stringless beans, the LFL is typically absent. In contrast to the clear anatomical differences between these three groups, no variation between PD-resistant and PD-susceptible domesticated dry bean pods was observed (Fig. 1B, 1C).

**Fig. 1.**
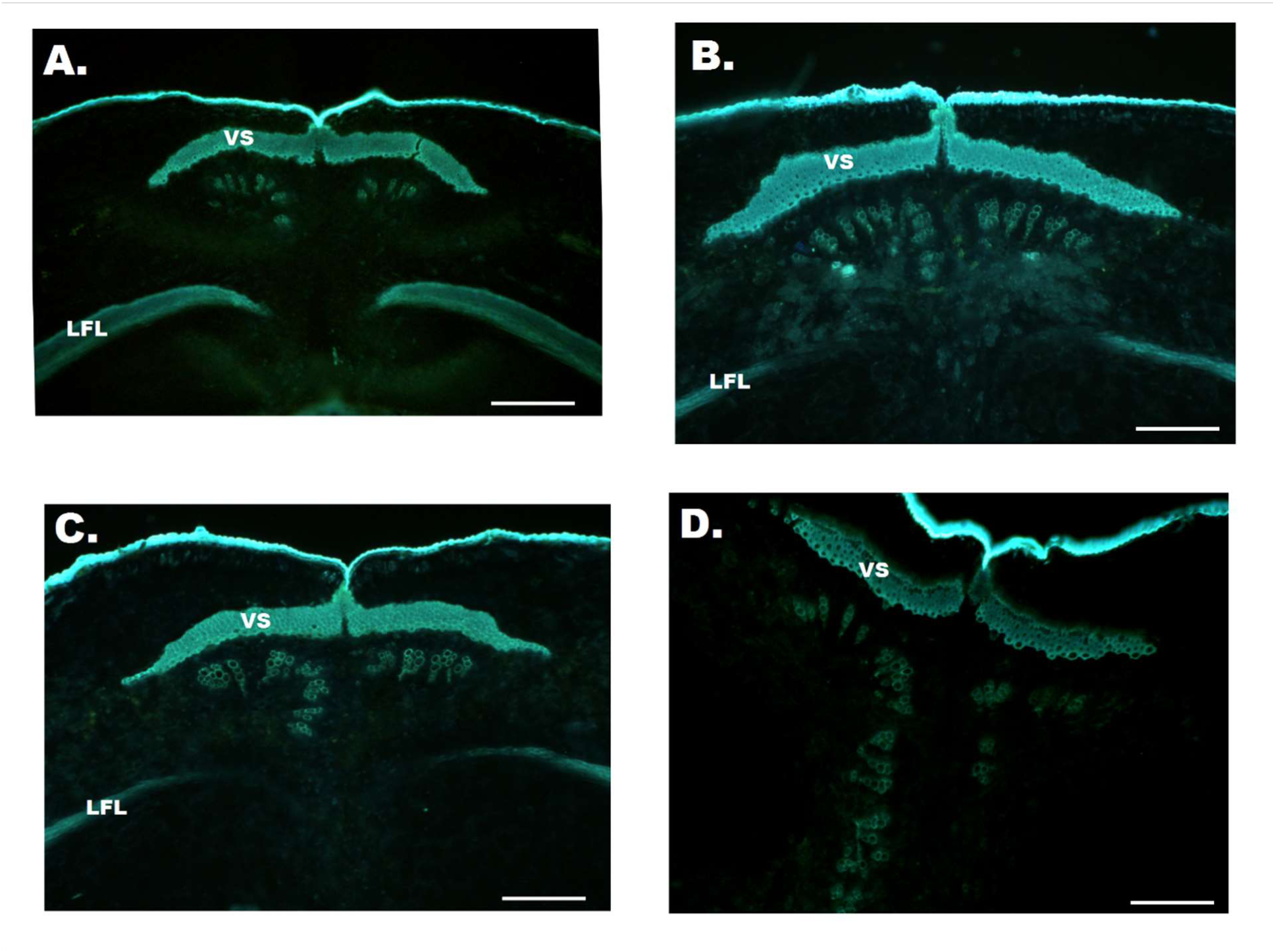
Variation in PD-related structures in common bean. (A) Cross-section of the ventral suture of G12873, a wild Middle American bean. Wild beans show very high pod dehiscence (PD) and extensive lignified vascular sheath (VS) and fiber layer (LFL) deposition in pod walls. (B) In PD-susceptible domesticated dry beans (cv. ICA Bunsi shown), LFL deposition is reduced relative to wild types, indicating that these cells may be related to Middle American common bean domestication. (C) PD-resistant dry beans (cv. SXB 405 shown) are anatomically similar to PD-susceptible domesticated types (see B). (D) Stringless varieties (cv. Midas shown) display a reduction in VS lignification, including a reduction in secondary cell wall thickening. The LFL is absent in these varieties. Scale bars represent 100μm.

### Variation in the ICA Bunsi/SXB 405 (IxS) population

Segregation for PD was determined in an RI population derived from PD-susceptible cv. ICA Bunsi’ and PD-resistant cv. ‘SXB 405’. Both parental genotypes belong to the Middle American gene pool. Three phenotyping approaches were used to evaluate PD (Fig. S1) and each had a unique distribution pattern (Fig. S2). These phenotypes were strongly correlated (Fig. S3). Varieties that dehisced in the field had higher rates of PD after desiccation at 65°C (two-tailed t-test, p=3.1* 10^−8^) and required lower levels of force to induce fracture at the sutures (two-tailed t-test, p=1.2* 10^-9^). Similarly, the proportion dehiscing in the desiccator and force required to cause PD were negatively correlated (r^2^ = 0.71 simple linear model, p < 2*10^-16^).

QTL mapping identified a major, PD-related QTL peak located in the same position on linkage group Pv03 using each of the three phenotyping methods (Fig. 2). The QTL mapped between SNP markers ss715639553 and ss715639323, which are separated by approximately 900 kb of physical distance (Table 1). Force measurement produced the most significant results (LOD score 53.3), followed by desiccation (LOD score 42.7), and field notes (LOD score 8.9). All methods produced results that were statistically significant based on 1000 randomized permutations of the data. The allele at the most significant SNP explained 17% of the variation in PD based on field notes, 59% of the variation based on desiccation, and 64% of the variation in force required for fracture in the population. Analyses to find additional PD QTLs failed to identify other regions of interest in the IxS population.

**Table 1.**
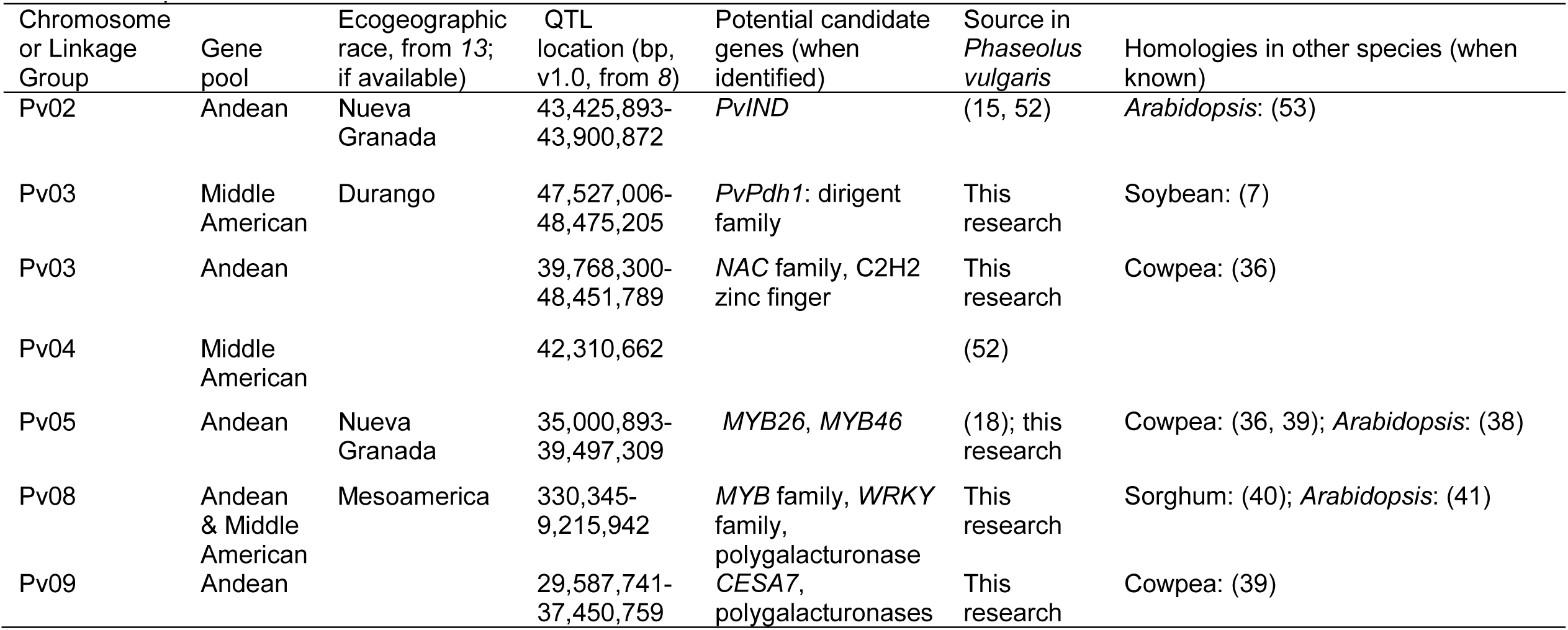
Summary of pod fiber or dehiscence QTLs, their genome locations, potential candidate genes, and homologies with other species

| Chromosome or Linkage Group | Gene pool | Ecogeographic race, from 13; if available) | QTL location (bp, v1.0, from 8) | Potential candidate genes (when identified) | Source in *Phaseolus vulgaris* | Homologies in other species (when known) |
| --- | --- | --- | --- | --- | --- | --- |
| Pv02 | Andean | Nueva Granada | 43,425,893-43,900,872 | *PvIND* | (15, 52) | *Arabidopsis*: (53) |
| Pv03 | Middle American | Durango | 47,527,006-48,475,205 | *PvPdh1*: dirigent family | This research | Soybean: (7) |
| Pv03 | Andean |  | 39,768,300-48,451,789 | *NAC* family, C2H2 zinc finger | This research | Cowpea: (36) |
| Pv04 | Middle American |  | 42,310,662 |  | (52) |  |
| Pv05 | Andean | Nueva Granada | 35,000,893-39,497,309 | *MYB26*, *MYB46* | (18); this research | Cowpea: (36, 39); *Arabidopsis*: (38) |
| Pv08 | Andean & Middle American | Mesoamerica | 330,345-9,215,942 | *MYB* family, *WRKY* family, polygalacturonase | This research | Sorghum: (40); *Arabidopsis*: (41) |
| Pv09 | Andean |  | 29,587,741-37,450,759 | *CESA7*, polygalacturonases | This research | Cowpea: (39) |

**Fig. 2.**
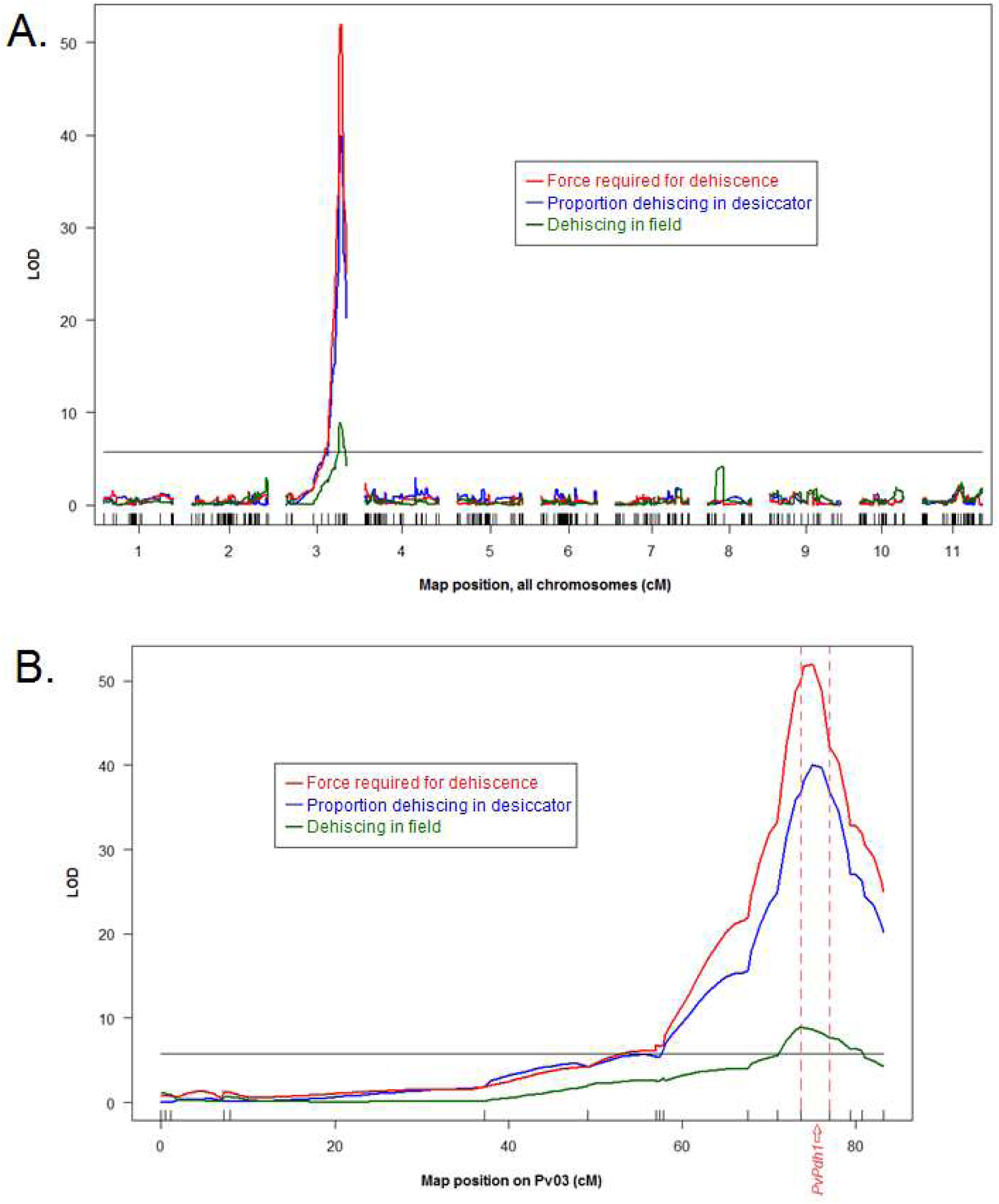
Pod dehiscence (PD) QTL mapping based on three phenotyping methods. (A) Genome-wide and (B) Pv03-specific mapping results. All methods produced statistically significant results in the same region of chromosome Pv03. The significance threshold, determined by 1000 randomized permutations of the data, is shown as a black bar at LOD=5.80. The common bean ortholog of *Pdh1*, which regulates PD in soybean, is located between the most significant markers from QTL mapping (Table S1).

### Synteny mapping and expression

Due to the close phylogenetic relationship and extensive microsynteny between *P. vulgaris* and *G. max* (19, 20), gene families known to affect PD in soybean were primary candidates for control of the trait in common bean. These families include the *NAC*-domain transcription factors and dirigent-like genes.

No *NAC*-domain transcription factors exist in the Pv03 QTL mapping interval. In contrast, the LegumeIP 2.0 synteny tool indicated that strong synteny exists between the soybean region surrounding *GmPdh1* and the common bean QTL (Table S3). This is in agreement with previous synteny analyses (20, 21). An amino acid BLAST of GmPDH1 (cv. Toyosume) against the *P. vulgaris* G19833 proteome (21) indicates that the most similar common bean protein is encoded by Phvul.003G252100, which maps between the two most significant Pv03 QTL SNP markers. A neighbor-joining tree of common bean and soybean dirigent proteins indicates that GmPDH1 and the protein product of Phvul.003G252100 cluster together (Fig. S4). Furthermore, the common bean gene’s expression is limited to developing pods, with no detectable expression in any other tissues (Fig. S5; data from (22)). This is comparable to the expression of soybean *PDH1* (7), and indicates that the gene serves a function unique to pods. Together, these results suggest that Phvul.003G252100 is orthologous to *GmPDH1.* Phvul.003G252100 is hereafter referred to as *PvPdh1.*

### Sequencing of PvPdh1

Sequencing of *PvPdh1* in ICA Bunsi and SXB 405 revealed a non-synonymous single-base-pair substitution at position 485 of the gene’s coding sequence (Fig. S6A). This substitution leads to a threonine/asparagine polymorphism (T162N) in the protein product (Fig. S6B). The 11 RILs with recombination between the most significant markers from QTL mapping showed complete co-segregation between the threonine/asparagine polymorphism and the PD phenotype (Table S1).

To investigate the functional importance of T162N, we evaluated the extent of its sequence conservation, surveyed literature related to this position in closely related dirigent proteins and used software tools to predict the effect of this substitution at the position. Sequencing of *PvPdh1* in several species of wild and domesticated *Phaseolus* from the USDA National Plant Germplasm System (NPGS) and UC Davis showed that the asparagine at this position was unique to the Middle American domesticated gene pool (Table S2). No polymorphism in the Andean gene pool was consistently associated with PD. In the Middle American gene pool, PD was significantly higher among genotypes with a threonine at position 162 than an asparagine (t-test: p=0.0002, n=47, Fig. S7). This threonine was strictly conserved in Andean domesticated common bean, Middle American and Andean wild common bean, and the closely related *P*. *lunatus* and *P*. *dumosus* (Table S2). The residue is present in 99 of the 100 most similar proteins in the NCBI database (Fig. S8A; see Discussion), indicating its functional importance. This threonine is also conserved in the 19 most similar proteins of *Selaginella moellendorffii* (Fig. S8B), a member of the first diverging group of lignin-containing plants. No comparable protein could be found in the proteome of *Physcomitrella patens*, a non-lignified moss.

Studies of closely related dirigent proteins indicate that this threonine is a component of one of the protein’s active sites (“T163” in Fig. S8C; from (23)), and its substitution eliminates protein function (Fig. S8D; from (24)). An analysis with PROVEAN (25) predicted that the T162N mutation would have a deleterious effect (score: −4.587, cutoff = −2.5).

### Validation through association mapping

The BeanCAP Middle American Diversity Panel (MDP) (26) was grown to determine, using the desiccation method, whether the Pv03 *PvPdh1* locus was related to PD in a broader population. A genome-wide association study (GWAS) indicated that the SNP closest to *PvPdh1* in physical distance was also the most significantly associated with PD (Fig. 3A, MAF threshold = 0.1). This SNP (S1_149243152) was 5.7 kb from the polymorphism in *PvPdh1.* Pv06 and Pv08 also included loci significantly associated with PD.

**Fig. 3.**
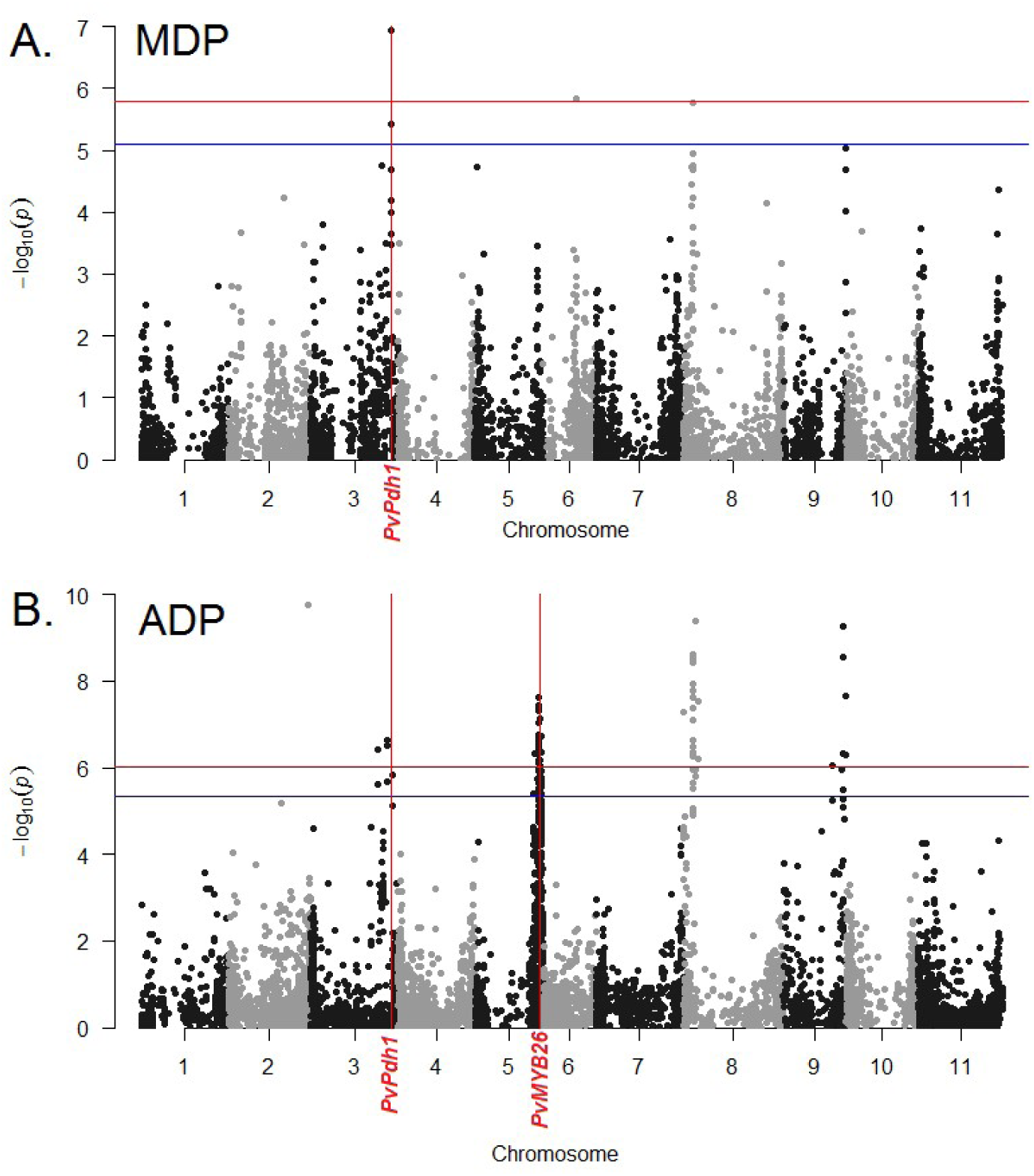
GWAS of PD in independently domesticated common bean populations. (A) In the Middle American Diversity Panel (MDP), the most significant SNP is located 5.7kbp from the *PvPdh1* putative causal polymorphism. Pv06 and Pv08 also included loci of interest. (B) In the Andean Diversity Panel (ADP), chromosomes Pv03, Pv05, Pv08, and Pv09 include major regions of interest. SNPs located near *PvMYB26* (18) on Pv05 were highly significant. Horizontal red and blue lines indicate the Bonferroni-corrected significance threshold for an alpha of 0.01 and 0.05, respectively. Based on the proportion of pods dehiscing in a desiccator, with correction for population structure by PCA.

GWAS was also conducted in the Andean diversity panel (ADP) (27) to determine which loci control PD in this independently domesticated population. Chromosomes Pv03, Pv05, Pv08, and Pv09 all included major regions significantly associated with PD (Fig. 3B). The QTL on chromosome Pv08 was in an overlapping physical position with a QTL from the MDP (Fig. 3A).

In both the Andean and Middle American gene pools, PD varied greatly between market classes (Table S4). In Andean beans, PD after desiccation averaged 3% in the purple speck market class and 41% in the cranberry types. In Middle American beans, averages were below 1% for pinto types of race Durango, and as high as 18% in the black beans of race Mesoamerica. Members of Middle American race Mesoamerica displayed considerable variation in PD. GWAS using only members of this race (MDP with PC1 > 50) showed that the Pv08 QTL was most closely associated with PD in the population (Fig. S9). SNP S1_329543689, near the center of this interval of interest, was used for subsequent analyses. The region near *PvPdh1* did not include significant SNPs in this race, further indicating that races Durango and Mesoamerica rely on different genes for PD resistance.

To visualize the correlation between PD and population substructure in the MDP, PD was plotted against the first principal component of the genetic data. Each point was color-coded by its allele at the GWAS SNP peaks on Pv03 (S1_l49243152, 5.7kb from *PvPdh1*) and Pv08 (SNP S1_329543689) (Fig. 4). Members of the MDP with the Pv03 PD resistance allele exhibited mean PD in the desiccator of 0.0067, with a maximum value of 0.14. In genotypes with no known resistance allele, the mean level of PD was 0.206 and ranged up to 0.63 (Fig. 4).

**Fig. 4.**
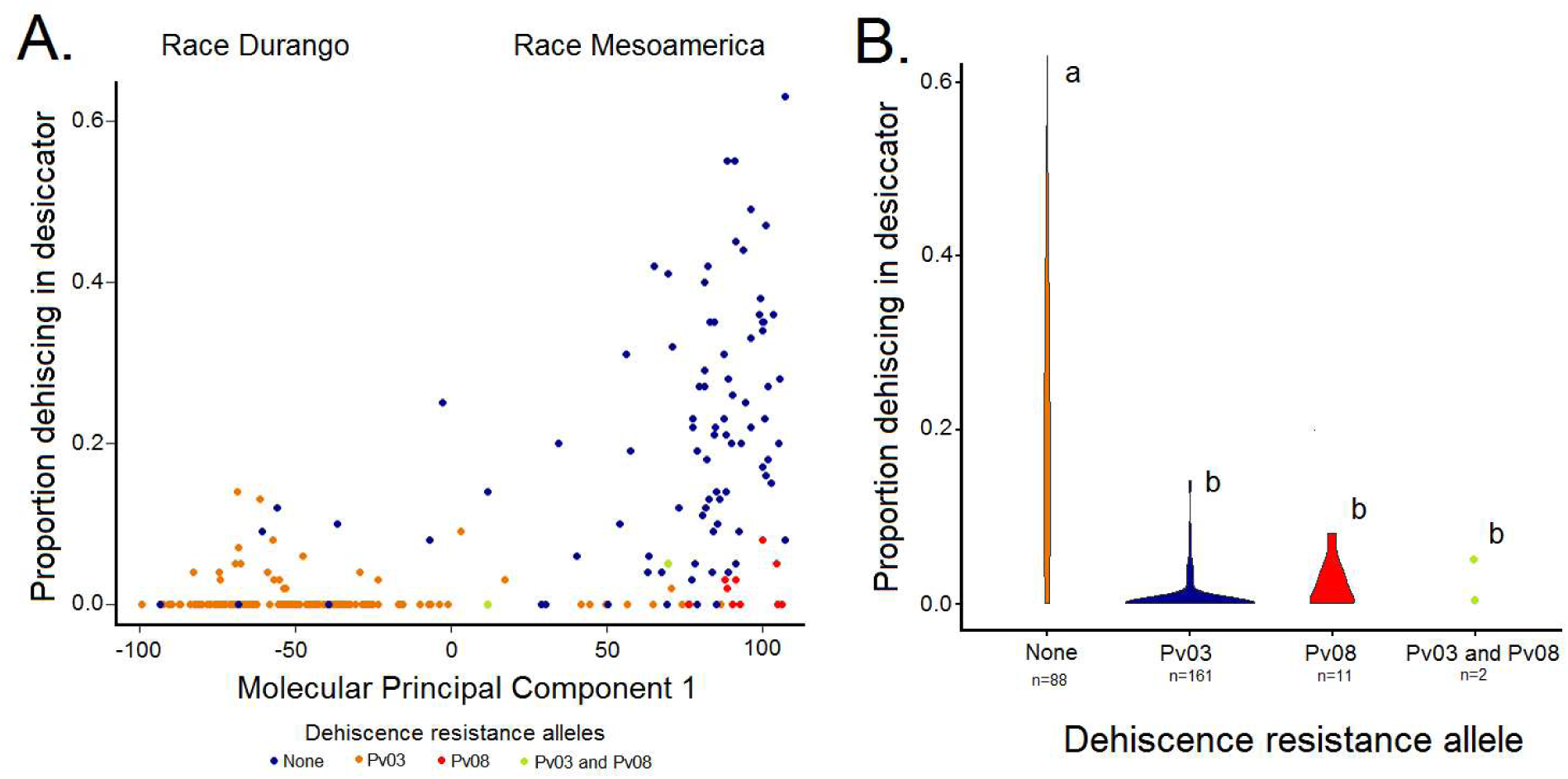
The relationship between PD, ecogeographic race, and resistance alleles. (A) The first principal component of genetic data for the MDP separates race Durango (at left) from race Mesoamerica (at right). Members of race Durango have low susceptibility to PD relative to members of race Mesoamerica. Accessions are color coded by genotype at the GWAS peaks on Pv03 and Pv08. (B) A violin plot showing of the extent of PD by allele in the MDP. Accessions with these PD resistance loci have significantly lower levels of PD than accessions with neither allele. Letters “a” and “b” distinguish significantly different groups (Tukey HSD).

## Discussion

A reduction in PD is a fundamental component of the domestication syndrome in common bean (15), and is important for future green and dry bean production and food security. In snap beans, a major gene - *St -* controls the presence or absence of pod strings and PD (Prakken, 1934; Koinange et al. 1996). For dry beans, we report here three novel QTLs mapped for the trait (on Pv03, Pv08, and Pv09), confirm a QTL on Pv05, identify a putative causal polymorphism in the *PvPdh1* gene underlying the major QTL on Pv03, and describe the association between Pv03, Pv05, and Pv08 QTLs and climate variables, especially precipitation.

### Anatomical differences among wild and domesticated types

Our microscopy results are consistent with those of previous researchers, who noted that reduced lignification in the LFL is correlated with reduced PD among wild beans, domesticated dry beans, and domesticated snap beans (2, 3, 28). However, we were not able to identify anatomical differences responsible for the variation in dehiscence among dry bean cultivars. This is consistent with the pattern seen in *GmPdh1* in soybean (8) and the expected result of a mutation in *PvPdh1.* Indeed, *PvPdh1* is thought to modify the biochemical structure of lignin, rather than its total quantity or cell fate in the relevant pod structures. A loss of function mutation in this gene would therefore not lead to clear anatomical differences relative to the wild-type.

### *PvPdh1* as a candidate gene for the Pv03 QTL identified in this study

The strict conservation of the threonine at position 162 in *PvPdh1* highlights its functional importance. Of the 100 most similar known protein models, the one that lacks a threonine at this position is found in *Trifolium subterraneum*, a legume that produces pods that mature underground. PD is not relevant for seed dispersal in this species, and the gene may be undergoing pseudogenization. This threonine is maintained in the proteome of *Selaginella moellendorffii*, indicating that the residue has been conserved since before the lycophyte-euphyllophyte divergence. This coincides with the origin of lignin and lignans, and indicates the residue’s functional role for members of the protein family. In a remarkable example of parallelism, independent loss of function mutations in this gene are found in certain domesticated populations in soybean (*G. max*) and in *P. vulgaris,* both species being subjected to selection for reduced dehiscence. This provides an additional example of strongly convergent phenotypic and molecular evolution (29). Similar examples of parallel evolution in common bean included the determinacy trait (*fin* or *PvTFL1y*; (30, 31)) and absence of pigmentation (*P*; (32)).

Further research is needed to identify the biochemical and biophysical aspects responsible for differences in PD in domesticated dry beans. Notably, our results could shed light on the fundamental process of lignin synthesis. Dirigent-like genes, including *PvPdh1,* encode non-enzymatic proteins that guide the dimerization of lignin and lignan monomers (33). The role of these proteins in lignin synthesis has been debated, with suggestions that polymerization is guided (34) or unguided (35). Varieties of common bean with mutations in *Pdh1* could be used to elucidate the role of this protein family in lignin synthesis generally.

### QTLs and Candidate Genes Identified by Association Mapping

Association mapping revealed several other dehiscence-related QTLs across the gene pools and races of common bean (Table 1). Our ADP association mapping identified significant Pv03 SNPs in an interval that is syntenic with a region controlling dehiscence in cowpea (36). *NAC* family and C2H2-type zinc finger transcription factors are found in this region (Table 1), and members of these families affect PD in soybean (6) and rapeseed (37), respectively. Orthologs of these genes may also affect dehiscence in cowpea (36). Interestingly, the QTL is large enough to include *PvPdh1*, although the QTLs discovered in Middle American beans and cowpeas are non-overlapping.

Another major QTL for PD in Andean beans maps to Pv05, as described recently (18), and several genes in this region are candidates for future study. Rau *et al.* (18) noted that an ortholog of *MYB26* exists in the qPD5.1-Pv region of interest on Pv05, which may be responsible for variation in PD. Significant Pv05 SNPs from our association mapping completely envelope the qPD5.1-Pv interval, supporting this result. Interestingly, our most significant Pv05 SNPs in the ADP are found just 22kb from *MYB46. MYB46* is involved in the same pathway as *MYB26* and the soybean PD resistance gene *SHAT1-5* (38, 6). *MYB46* also works redundantly with *MYB83*, a gene that may play a role in cowpea pod development (38, 39), making *MYB46* another potential subject of future study.

Several genes of interest exist near the middle of the ADP’s Pv08 GWAS peak. These include a MYB family transcription factor with similarity to *A. thaliana MYB17,* three *WRKY* family transcription factors, which are related to genes involved in sorghum dehiscence (40), and a polygalacturonase, a group known to influence PD in *A. thaliana* (41) (Table 1).

The Pv09 GWAS peak found in the ADP included a gene predicted to be *cellulose synthase A7* (*CESA7*, Table 1). *CESA7* may play a role in fiber development in cowpea (39). Similarly, two polygalacturonases are found in this interval, and members of this family are known to affect seed dispersal in *A. thaliana* (41). These genes may regulate dehiscence by altering the breakdown of cell wall material in developing pods. Identifying polymorphisms in PD candidate genes will be a promising area for future study.

### Associations with environmental conditions

PD in common bean is correlated with environmental parameters. The PD-resistant allele of *PvPdh1* on Pv03 is found exclusively in genotypes with ancestry from race Durango (Table 1). Race Durango is adapted to higher elevations and lower humidity regions, particularly in the northern part of Mexico (13). The semi-arid conditions in these areas cause pods to become dry and brittle, which exacerbates PD. The non-functional *PvPdh1* allele rose to very high frequency in this ecogeographic race. In contrast, race Mesoamerica is adapted to humid lowlands, where environmental conditions mask PD and reduce selection pressure against it. In this race, the loss-of-function *PvPdh1* allele remains at low frequency and PD is widespread (Fig. 4A). Interestingly, this ecogeographic pattern closely parallels that of soybean, in which *Pdh1*-mediated resistance to PD is most common in arid regions (7). *PvPdh1* may also be responsible for the ease of threshing that has been noted in race Mesoamerica (13). In humid environments, the wild type *PvPdh1* allele may facilitate separation of seeds from pod material, while PD in the field remains low. In northern Mexico, the semi-arid climate facilitates threshing but increases PD in the field. Under these conditions, the PD-resistant allele is advantageous. Because of this trade-off, the polymorphism in *PvPdh1* appears to be related to local adaptation (Fig. 5). Alleles that prevent PD will be valuable in coming decades, which are predicted to be increasingly arid (42).

**Fig. 5.**
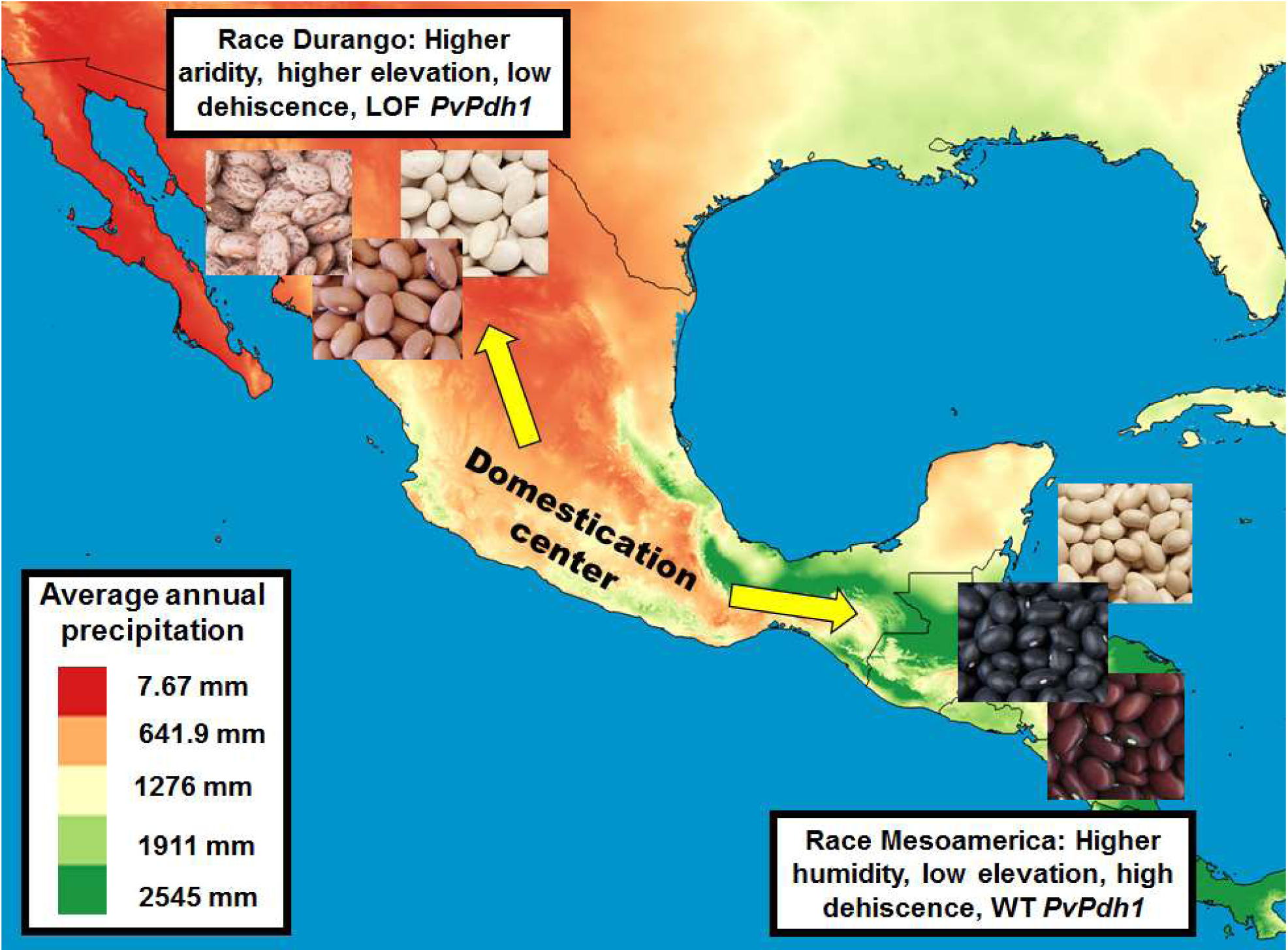
*PvPdh1* is related to local adaptation and range expansion in common bean. PD is nearly absent in Race Durango, a group adapted to the hot, dry environments of northern Mexico, where environmental aridity exacerbates PD. The loss of function *PvPdh1* allele is nearly at fixation in this population. In contrast, race Mesoamerica is adapted to humid lowlands, where conditions mask PD susceptibility. PD has been selected against less strongly in this population, and the wild type *PvPdh1* predominates. For detailed information on the geographic distribution of these races, see Singh *et al.* (13).

### Redundancies in genetic control and maintenance of atavistic traits

Crosses between races have tremendous potential for crop improvement (for example, between races Durango and Mesoamerica (43)), but could also result in problematic gene complementation. Because several genes influence PD redundantly, cultivars descended from crosses between races could demonstrate atavistic transgressive segregation. This may be responsible for the high levels of dehiscence seen in some varieties of common bean. The interactions between these loci will also be of considerable importance for plant breeders.

## Methods

Details regarding materials and methods can be found in SI Methods. Pods were sectioned for microscopy using a Vibratome. Lignified and hydrophobic structures were visualized using epifluorescence microscopy after staining with Auramine O.

The IxS population was genotyped using the Illumina Infinium II BARCBean6K_3 BeadChip. In Spring 2014, 238 RILs were grown in an unreplicated trial and visually phenotyped for the presence or absence of PD. In fall 2014, 191 RILs in a partially replicated trial were phenotyped based on proportion of pods dehiscing due to desiccation and force required to cause fracture. The maximum LOD score of 1000 random permutations of the data was used as a significance threshold.

Synteny mapping was conducted using LegumeIP 2.0 (44) and CoGe SynMap (45). Candidate genes were identified by NCBI BLAST and clustered through the NCBI portal. Gene expression data were accessed through the Common Bean Gene Expression Atlas (22). *PvPdh1* of ICA Bunsi, SXB 405, and RILs of interest was amplified by PCR and sequenced at the UC DNA Sequencing Facility. The COnstraint-Based multiple ALignment Tool (COBALT) (46) was used to align the PvPDH1 amino acid sequence to the most similar documented proteins of the NCBI database. PROVEAN (25) was used to estimate mutational effects.

*PvPdh1* was sequenced in accessions with known pod shattering phenotypes from NPGS and UC Davis. Because members of the reproductively isolated Andean gene pool did not carry the T162N substitution, these individuals were filtered from subsequent analyses. For Middle American accessions, the categorical shattering scale used by USDA was translated into a simple numeric scale and PD between allele groups was compared using a student’s t-test.

The ADP (27) consisted of 208 phenotyped accessions, and these were evaluated based on presence of PD in the field, proportion dehiscing in the desiccator, and force required to cause fracture. The MDP (26) included 278 phenotyped varieties that were evaluated by the desiccation method alone. GWAS in both populations were conducted using TASSEL (47) through SNiPlay (48). Manhattan plots were visualized using the qqman R package (49).

Worldclim2 precipitation data (50) were compiled with Natural Earth national boundary shapefiles (51) in QGIS to visualize precipitation patterns in the range of Middle American beans.

## ACKNOWLEDGEMENTS

The authors would like to thank S. Beebe (CIAT, Cali, Colombia) for providing seeds of the IxS population. Seeds of the ADP and MDP were generously provided by R. Lee and P. McClean (North Dakota State University). Undergraduates M. Rocha, P. Silva Rezende, G. Coelho Portilho, N. Hamada, E. Yang, A. Herrera, J. Pimentel, M. Bustamante, E. White, J. Gonzales, and P. Augello contributed to DNA extractions, pod phenotyping, and other lab protocols. Paola Hurtado, Andrea Ariani, and other members of the Gepts lab provided ideas for data analysis. Funding for T.A.P. was provided through a Clif Bar Family Foundation Seed Matters fellowship and Lundberg Family Farms research support.

## Supporting Information

### SI Methods

#### Microscopy

Pods of G12873 (wild, high dehiscence), ICA Bunsi (domesticated dry bean, dehiscence susceptible) SXB 405 (domesticated dry bean, dehiscence resistant), and Midas (domesticated snap bean, dehiscence susceptible) were Vibratome-sectioned to identify morphological differences that might be associated with PD. All sectioned pods were greenhouse-grown and harvested when pods were at full size with seeds filled, at the onset of pod color change. All sections were 100 micrometers thick and made in a transverse plane perpendicular to the fibers of interest. All sections were treated with Auramine O for at least 20 minutes. Fluorescence was visualized using an Olympus microscope.

#### RI population and phenotyping for pod dehiscence

A recombinant inbred (RI) population developed from a cross between ICA Bunsi (domesticated, PD-susceptible) and SXB 405 (domesticated, PD-resistant) was used for QTL mapping (1). The population (IxS) of 238 RILs was field-grown during the spring and summer of 2014. The spring planting was an un-replicated trial conducted in Coachella, California. At maturity, plots were visually evaluated for the presence or absence of PD, and the data were used as a phenotype for QTL mapping. During the summer of 2014, the RI population was grown in a replicated trial in Davis, California. At maturity, dried pods from 191 RILs were harvested from each plot; these were evaluated for susceptibility to PD by two methods. First, all pods were desiccated at 65°C for seven days, and then returned to room temperature for a minimum of seven additional days. The proportion of pods dehiscence in this process was recorded for each plot. Second, the amount of force required to induce pod fracture was measured using an Imada force measurement gauge (method modified from (2)). A bit mounted to the gauge was used to press the ventral side of each pod at the most apical seed, and the peak force required to cause fracture at the apical end of the pod beak was recorded. Force required for PD was normalized to account for small but significant differences between note-takers, and the standardized score was used for QTL mapping.

#### Genotyping

Genomic DNA was extracted from parents and RILs of the IxS population using a modified CTAB protocol. DNA quality was confirmed using a NanoDrop spectrophotometer. The IxS population was genotyped using the Illumina Infinium II BARCBean6K_3 BeadChip (3); 382 segregating SNPs were identified in the population. Primers spanning the transcribed sequence of Phvul.003G252100, a candidate gene underlying the major QTL identified in this study, were developed using the NCBI Primer-BLAST tool. Differences in the genomic sequence around *PvPDHI* exist between the Middle American and Andean gene pools, so variable PCR primers were used between the gene pools. PvPdh1ALL MA Forward (CATCTCCCCCATTTTCCCCC) and PvPdh1ALL Reverse (AACACGTGGAAGAGGAGGATT) were used for Middle American accessions, while PvPdh1ALL Andean Forward (CATCTCTCCCATTTTCTCCT) and PvPdh1ALL Reverse (AACACGTGGAAGAGGAGGATT) were used for Andean types. PCR conditions for this amplification included an initial denaturation at 95°C for 180s, 38 cycles of 95°C for 30s, 51 °C for 30s, and 68°C for 60s, and a final elongation step of 68°C for 300s. PCR products were cleaned using a GeneJET PCR Purification Kit and sequenced at the UC DNA Sequencing Facility by Sanger sequencing.

#### QTL mapping

Composite interval mapping was conducted using the R package R/qtl (4). Field dehiscence, proportion dehiscing in a desiccator, and force measurements were separately used to identify PD QTLs marked by SNPs. The maximum LOD score of 1000 randomized permutations of the data was used as a significance threshold. Multiple QTL mapping was conducted using the scantwo function in R/qtl and by running the analysis with RILs subsetted by genotype at the most significant marker near *PvPdh1* on Pv03.

#### Synteny mapping and expression

Candidate genes related to PD were identified in Phytozome 12 (5). Synteny comparisons between common bean and soybean were made using the Legume Information System 2.0 (6); these were verified using available literature (7, 8). The CoGe SynMap (9) and LegumeIP 2.0 (6) synteny tools were used to compare syntenic regions between *Arabidopsis* (Col-0, TAIR10), common bean (G19833, Pvulgaris_V1.0_218; (8)), and soybean (Williams 82, Release 1.1; (10)). For tree generation, the PvPDH1 amino acid sequence was BLASTed against the *A. thaliana, G. max*, and *P. vulgaris* proteomes. Default Grishin settings were used to construct the distance matrix. A fast-minimum evolution tree (11) was generated based on a maximum sequence difference of 0.85. Gene expression from a variety of tissues and developmental stages were based on published data (12) and visualized in R.

#### Amino acid conservation analyses

The complete amino acid sequence of *PvPdh1* from accession G19833 was BLASTed against the NCBI proteome database. The COnstraint-Based multiple ALignment Tool (COBALT) (13) was used to align the most similar proteins known among several plant taxa and identify conserved residues. The Protein Variation Effect Analyzer (PROVEAN) software tool (14) was used to estimate the effect of mutations of interest.

#### Validation of the role of *PvPdh1* in a wider population

The Genetic Resources Information Network (GRIN) database of the National Plant Germplasm (NPGS) includes PD phenotype data for the genus *Phaseolus.* PD-susceptible and PD-resistant varieties from this pool were selected for validation of the role of *PvPdh1* in PD. A small number of varieties commonly grown at UC Davis with known PD phenotypes were also genotyped. Stringless snap bean varieties were specifically excluded from the analysis to avoid the epistatic effect of the *Stringless (St)* locus on PD. Genomic DNA was extracted using a modified CTAB method; amplification and Sanger sequencing of *PvPdh1* were conducted as described previously. Genotypes were separated into the Andean or Middle American gene pool based on an indel in the 3’ UTR of *PvPdh1.* This indel consistently predicted the gene pool in varieties of known ancestry. After sequencing, Middle American varieties were divided into groups based on amino acid at position 162 of *PvPdh1.* The degree of dehiscence between these groups was evaluated by student’s t-test.

#### Validation of QTL mapping results using association mapping

Two hundred and eight accessions of the Andean Diversity Panel (ADP) (15) were grown in Davis, CA during summer 2016. PD in the field, proportion dehiscing in a desiccator, and force required for fracture were recorded. Principal component analysis was conducted on SNP data for the population, and the results were used as covariates to account for population structure. Two hundred seventy-eight members of the BeanCAP Middle American Diversity Panel (MDP) (16) were phenotyped for PD by desiccation in 2017. Association mapping was conducted using GLM in TASSEL (17) via SNiPlay (18). All results were visualized using the qqman R package (19).

#### Precipitation map generation

Precipitation across the native range of Middle American beans was mapped in QGIS 2.18.19 using data from worldclim2 (20). National boundaries and coastlines were added using shapefiles available through Natural Earth (21). USGS topographical global raster data grids were also used to improve the visualization of coastlines (https://topotools.cr.usqs.gov/qmtedviewer/gmted2010globalgrids.php).

## Supplemental Figures

**Fig. S1.**
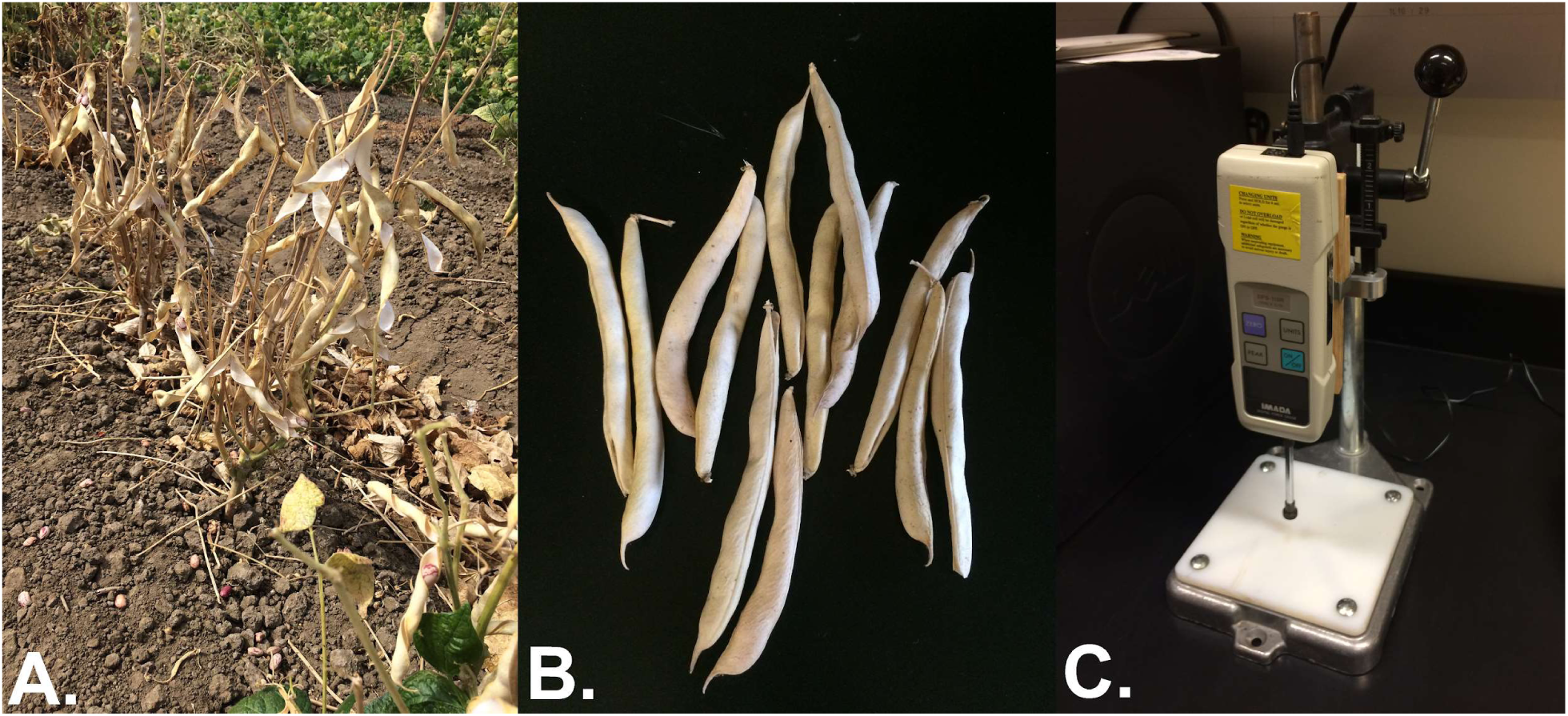
Phenotyping methods. PD was evaluated by (A) visual inspection of PD in the field, (B) proportion of dehiscing pods in a desiccator (none dehiscing in this sample), and (C) force required to induce fracture with a force measurement gauge.

**Fig. S2.**
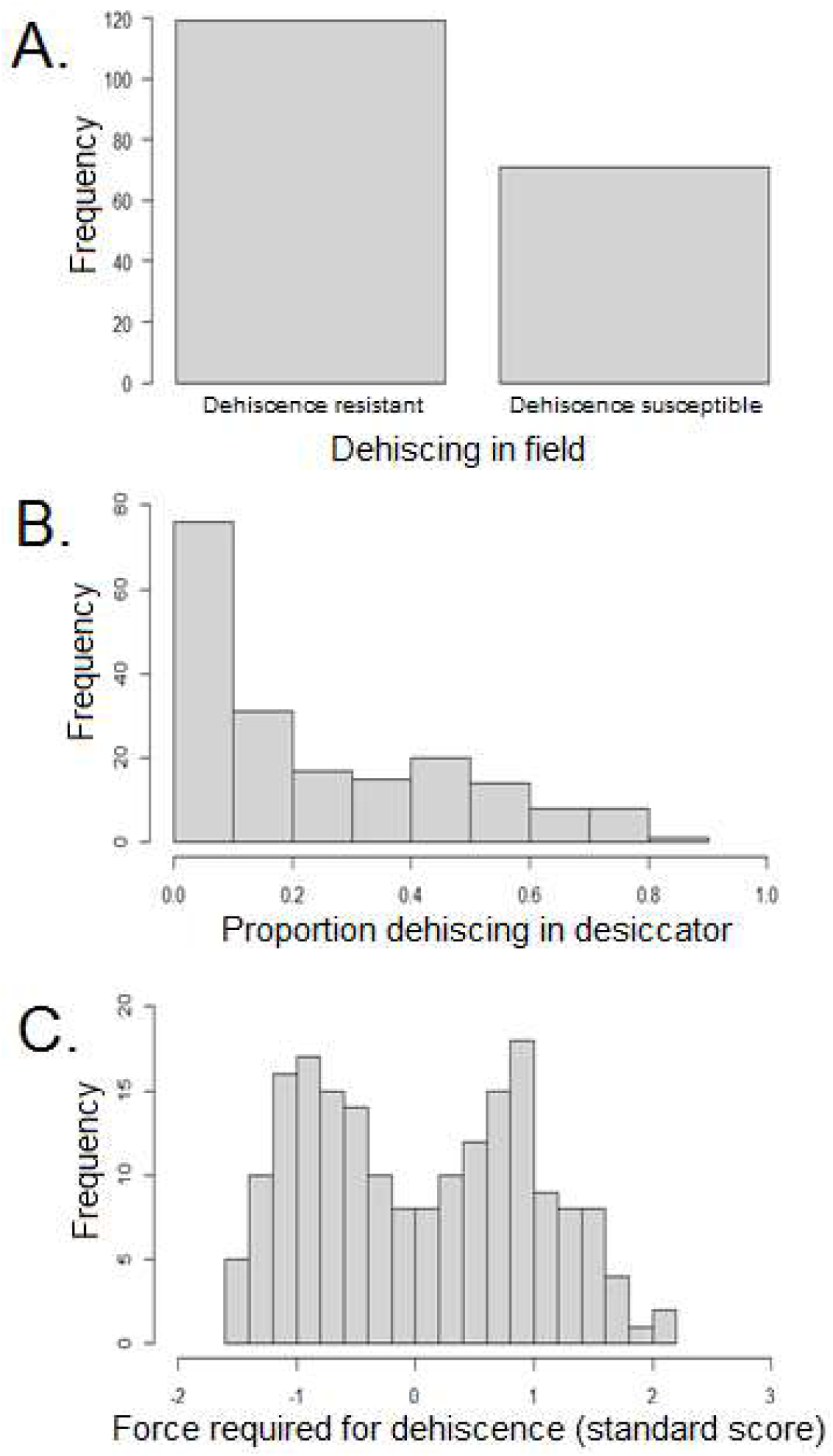
Phenotyping distributions in the ICA Bunsi/SXB 405 RI population. (A) Presence/absence of PD in the field. (B) The proportion of pods dehiscing after desiccation. C) The force required for pod fracture. Force measurements resulted in a bimodal distribution, indicating that a single large-effect gene was responsible for much of the population’s variation. “Frequency” represents the number of RILs falling into each bin.

**Fig. S3.**
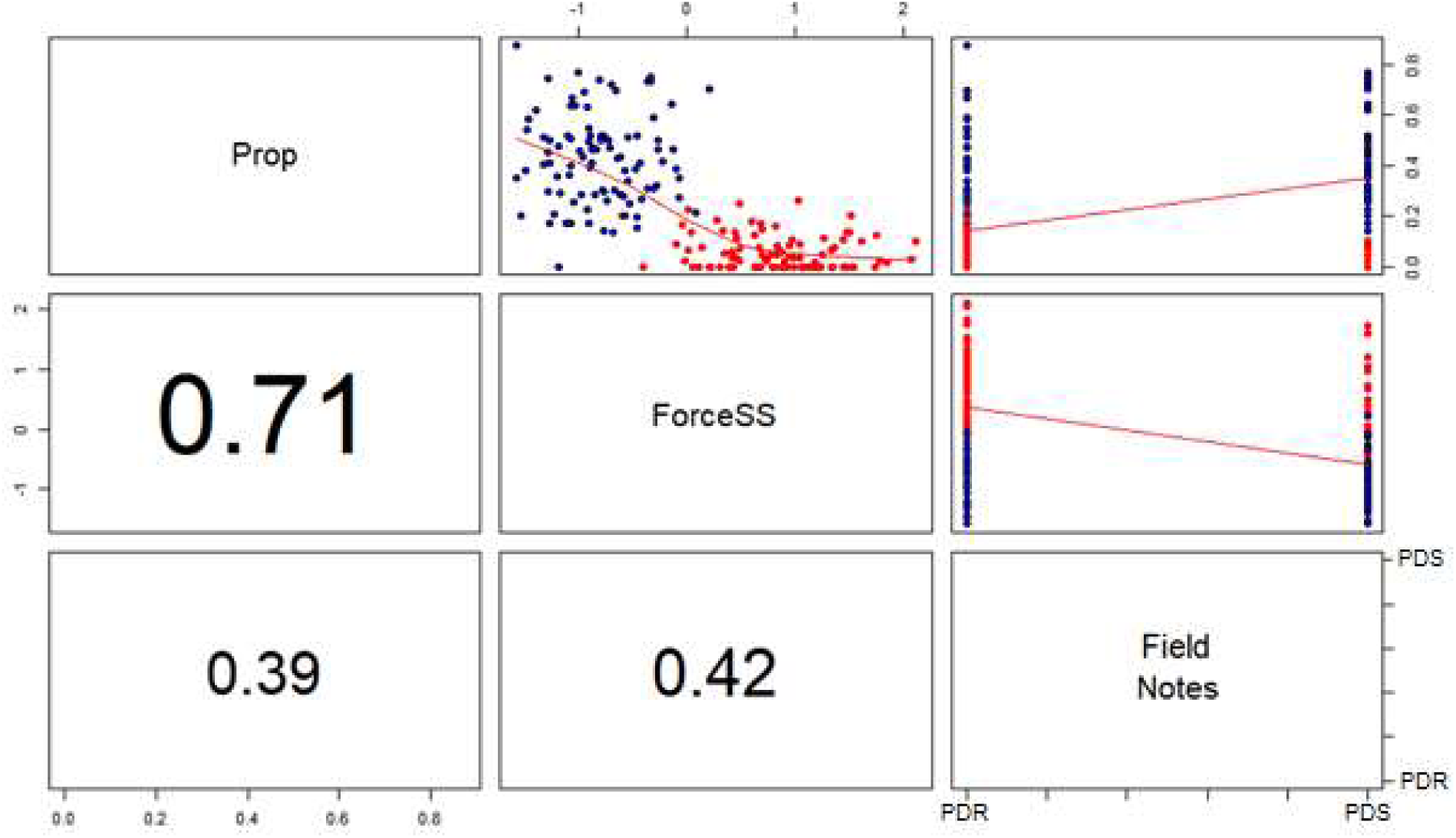
Correlations between phenotyping methods in the IxS RI population. RI lines are color coded by genotype at the *PvPdh1* locus. The numbers in the lower left panels indicate the correlation coefficients between those methods. PDS=PD susceptible, PDR=PD resistant.

**Fig. S4.**
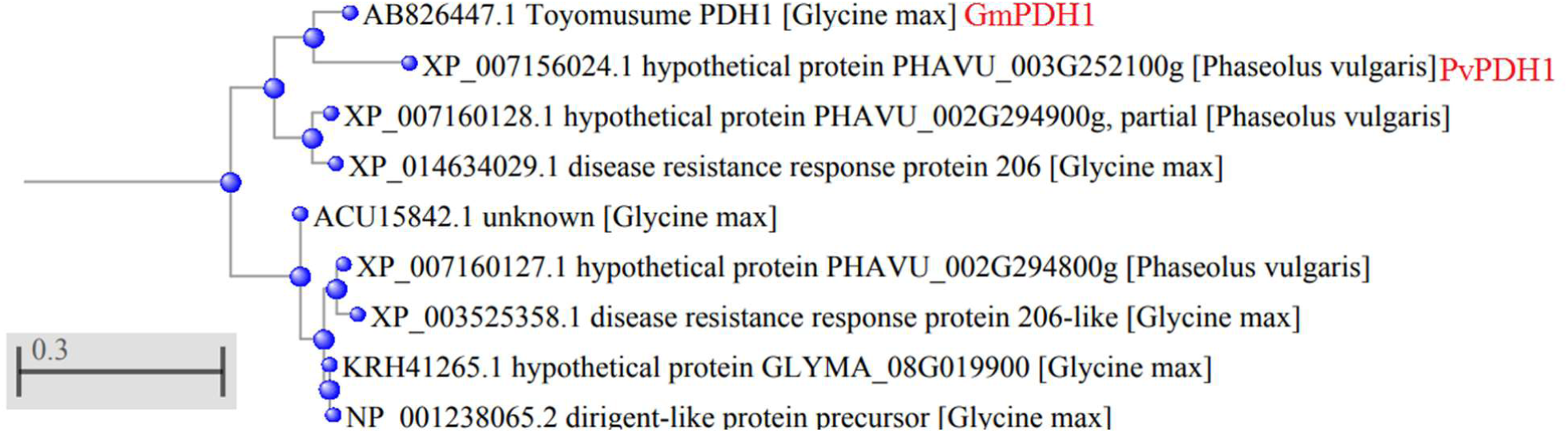
A rooted neighbor joining tree based on sequence of GmPDH1, PHAVU_003G252100g, and the most similar dirigent proteins of *G. max* and *P. vulgaris* in the NCBI database. GmPDH1 and PHAVU_003G252100g form a clade among all the proteins of these species, supporting their orthology. Tree derived from a Grishin protein distance matrix and rooted using 12 distantly related dirigent-like proteins.

**Fig. S5.**
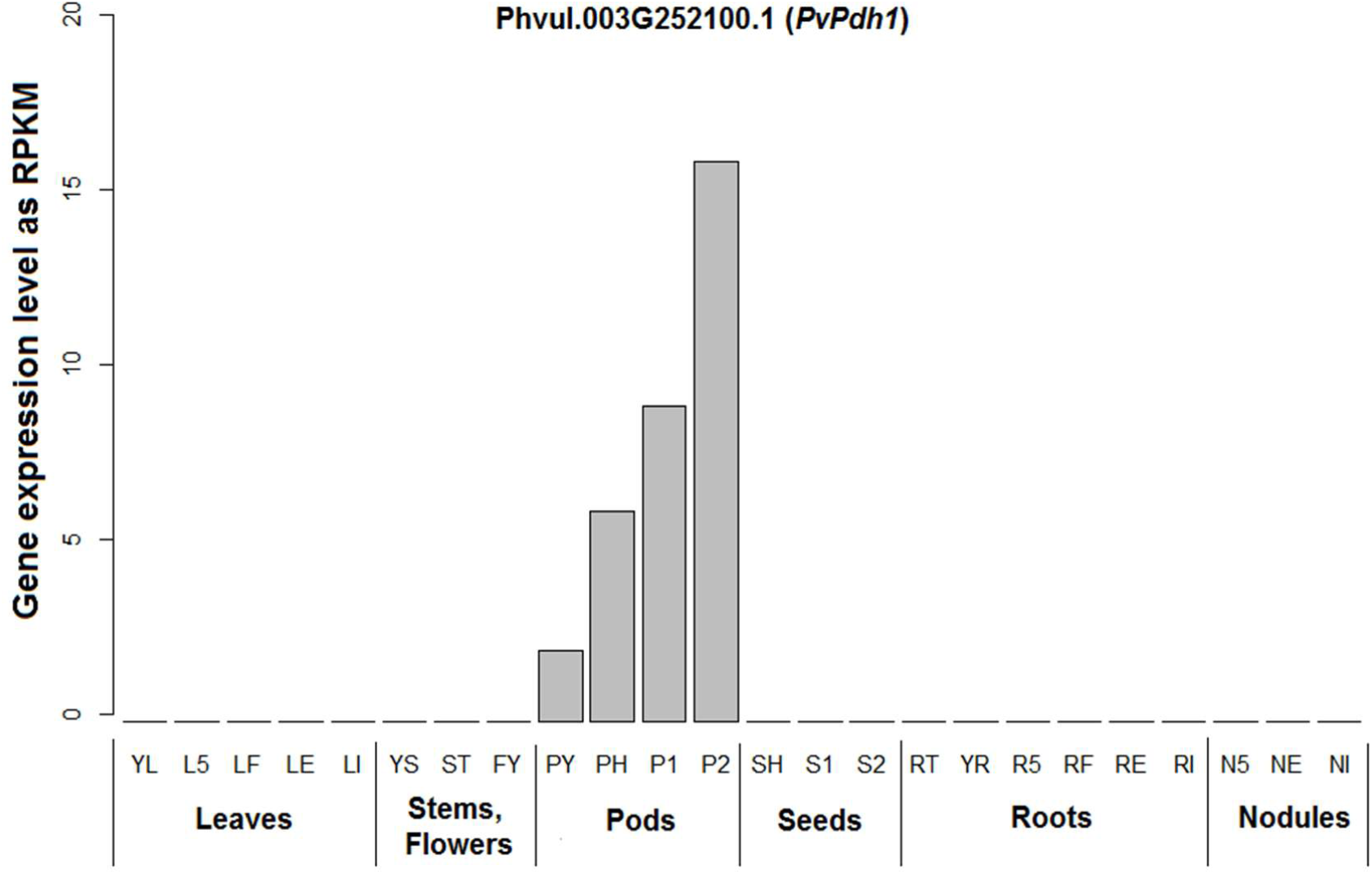
Expression of Phvul.003G252100.1 (*PvPdh1*) is unique to pods in *P*. *vulgaris* cv. Negro Jamapa. This pattern is extremely unusual even among homologs of *Arabidopsis* genes affecting PD. YL-Fully expanded 2nd trifoliate leaf tissue from fertilized plants L5-Leaf tissue collected 5 days after plants were inoculated with effective rhizobium; LF-Leaf tissue from fertilized plants collected at the same time of LE and LI; LE-Leaf tissue collected 21 days after plants were inoculated with effective rhizobium LI-Leaf tissue collected 21 days after plants were inoculated with ineffective rhizobium; YS-All stem internodes above the cotyledon collected at the 2nd trifoliate stage; ST-Shoot tip, including the apical meristem, collected at the 2nd trifoliate stage; FY-Young flowers, collected prior to floral emergence; PY-Young pods, collected 1 to 4 days after floral senescence. Samples contain developing embryos at globular stage PH-Pods approximately 9cm long, associated with seeds at heart stage (pod only); P1-Pods between 10 and 11 cm long, associated with stage 1 seeds (pod only); P2-Pods between 12 and 13 cm long associated with stage 2 seeds (pod only); SH-Heart stage seeds, between 3 and 4 mm across and approximately 7 mg S1-Stage 1 seeds, between 6 and 7 mm across and approximately 50 mg; S2-Stage 2 seeds, between 8 and 10 mm across and between 140 and 150 mg; RT-Root tips, 0.5 cm of tissue, collected from fertilized plants at 2nd trifoliate stage of development; YR-Whole roots, including root tips, collected at the 2nd trifoliate stage of development; R5-Whole roots separated from 5 day old pre-fixing nodules; RF-Whole roots from fertilized plants collected at the same time as RE and RI; RE-Whole roots separated from fix+ nodules collected 21 days after inoculation; RI-Whole roots separated from fix-nodules collected 21 days after inoculation; N5-Pre-fixing (effective) nodules collected 5 days after inoculation; NE-Effectively fixing nodules collected 21 days after inoculation; NI-Ineffectively fixing nodules collected 21 days after inoculation. From O’Rourke *et al.* (12).

**Fig. S6.**
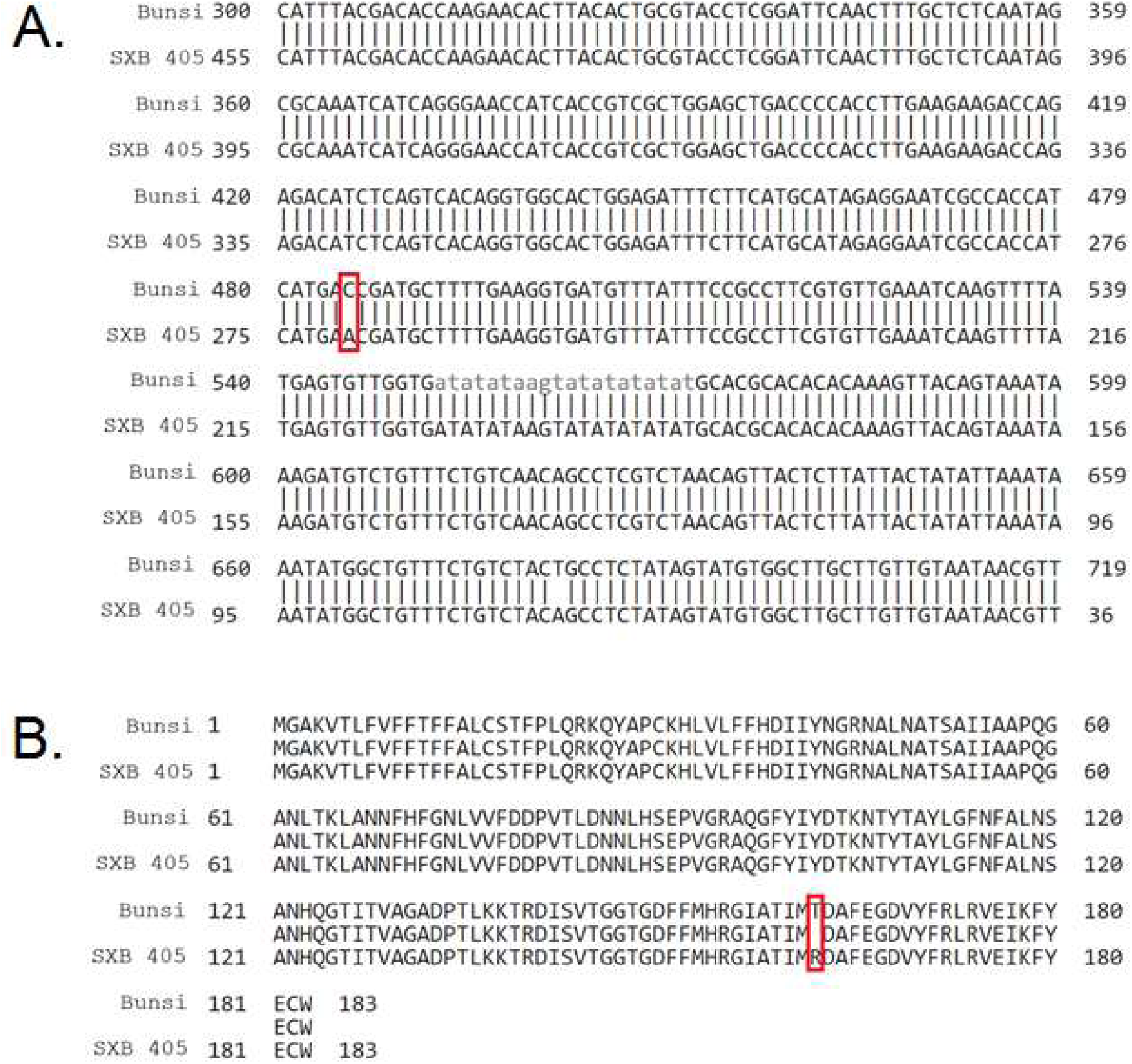
A polymorphism exists in *PvPdh1* between the parents of the RI population. A) At position 485 of the CDS of *PvPdh1,* there is a C/A polymorphism between ICA Bunsi and SXB 405. This nonsynonymous substitution leads to B) a threonine/asparagine polymorphism at position 162 in the amino acid sequence of the protein products.

**Fig. S7.**
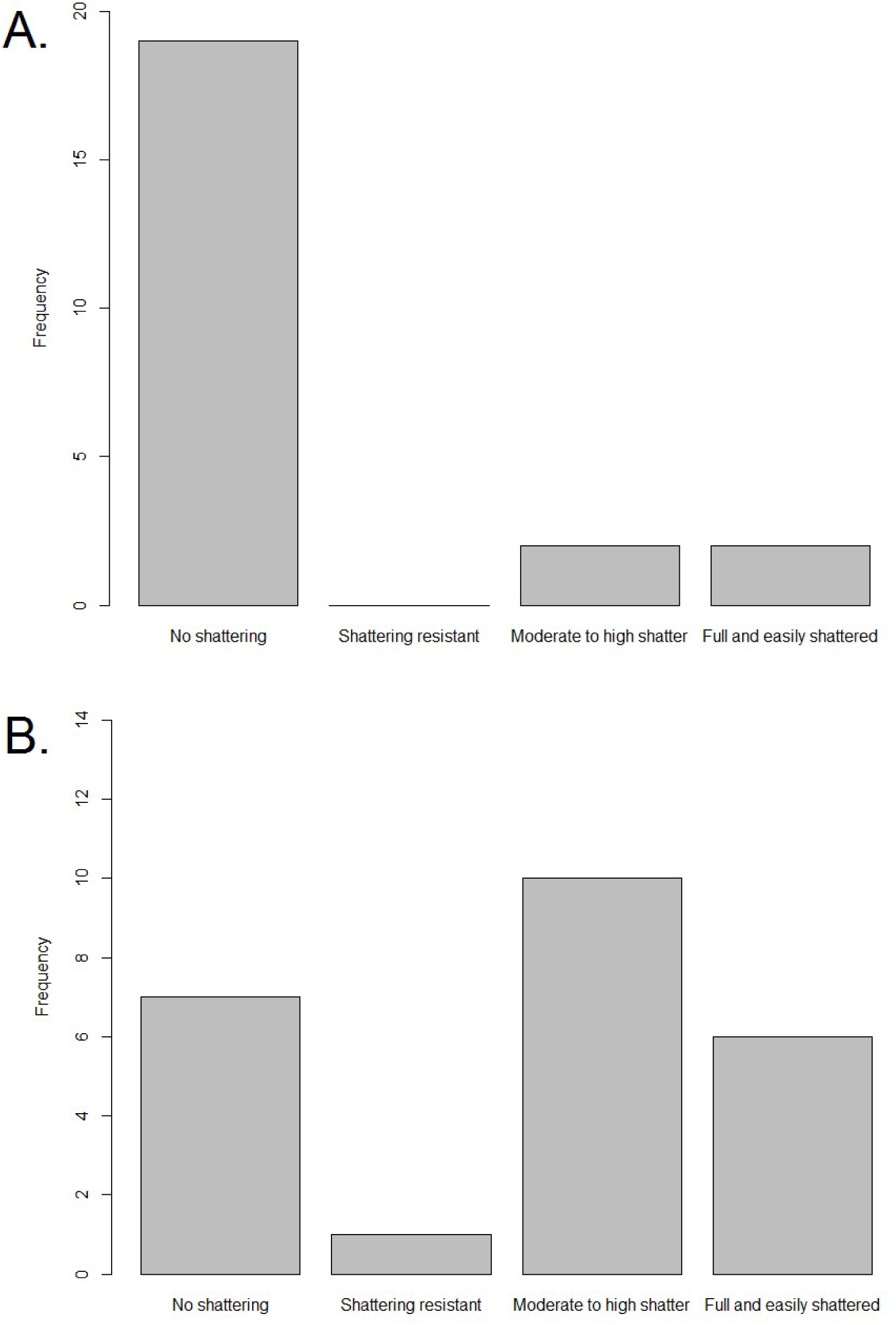
Dehiscence in Middle American GRIN NPGS accessions. A) In individuals with an asparagine at position 162, dehiscence resistance predominates. B) In individuals with a wild-type threonine at position 162, dehiscence susceptibility predominates. Accessions were phenotyped by GRIN NPGS and genotyped by Sanger sequencing of *PvPdh1.*

**Fig. S8.**
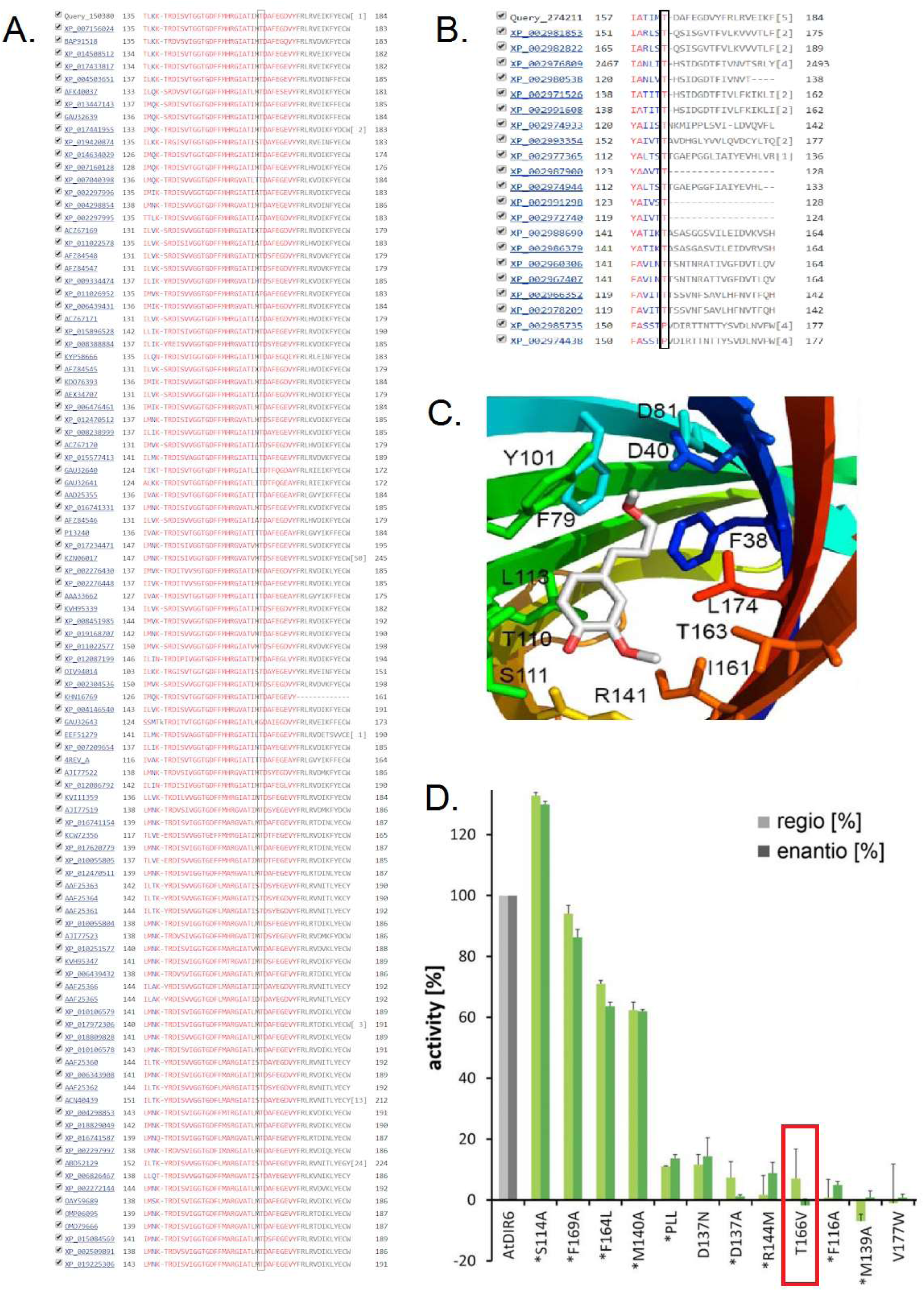
The threonine at position 162 is a highly conserved component of the active site for dirigent-like genes. (A) Of the 100 most similar proteins to PvPDH1 in the NCBI database, 99 have a threonine at the aligned position, indicating it is vital for protein functionality. The one exception is a gene from *Trifolium subterraneum*, which places pods underground and the gene may be undergoing gene decay. (B) The 19 most similar dirigent-like genes from *Selaginella moellendorffii* have a threonine at this position, indicating that the residue has been very strongly conserved for over 400 million years (22, 23). (C) In the closely related protein PsDIR6, the homologous threonine (T163) is an important component of the active site (from (24)). (D) Targeted mutagenesis of the equivalent residue (T166) in a closely related *Arabidopsis* protein showed that substituting the threonine with a valine led to a complete loss of gene function (from (25)).

**Fig. S9.**
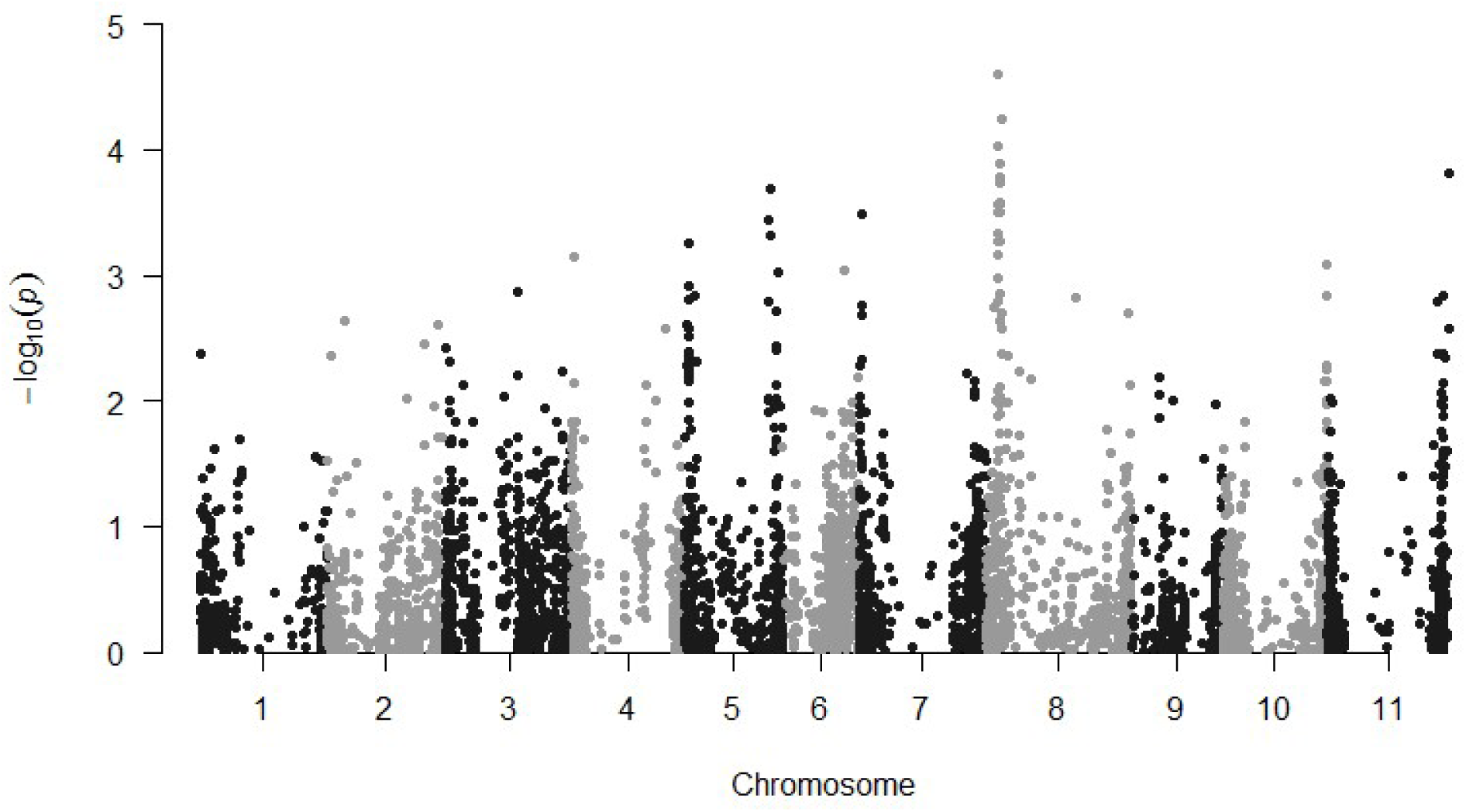
GWAS of pod dehiscence (PD) in Race Mesoamerica (MDP, PC1>50) using GLM in SNiPlay/TASSEL. Pv08 was most significantly associated with variation in PD, although no SNPs achieved significance in this smaller population, the most significant SNPs were located in an overlapping interval on Pv08 as a major QTL of the ADP, indicating that the same gene may be responsible for the variation across populations. MAF threshold = 0.1.

## Supplemental Tables

**Table S1.** Co-segregation between dehiscence phenotype and position 162 in *PvPdh1.* The 11 RILs with recombination between the flanking markers from QTL mapping showed perfect correspondence between phenotype and genotype at this position.

| RIL | Phenotype | *PvPdh1* CDS residue<br>485 | PvPDH1 peptide residue 162 |
| --- | --- | --- | --- |
| 013 | Dehiscent | Cytosine | Threonine |
| 036 | Dehiscent | Cytosine | Threonine |
| 052 | Dehiscent | Cytosine | Threonine |
| 094 | Dehiscent | Cytosine | Threonine |
| 096 | Dehiscent | Cytosine | Threonine |
| 006 | Non-dehiscent | Adenine | Asparagine |
| 033 | Non-dehiscent | Adenine | Asparagine |
| 065 | Non-dehiscent | Adenine | Asparagine |
| 101 | Non-dehiscent | Adenine | Asparagine |
| 117 | Non-dehiscent | Adenine | Asparagine |
| 154 | Non-dehiscent | Adenine | Asparagine |

**Table S2.**
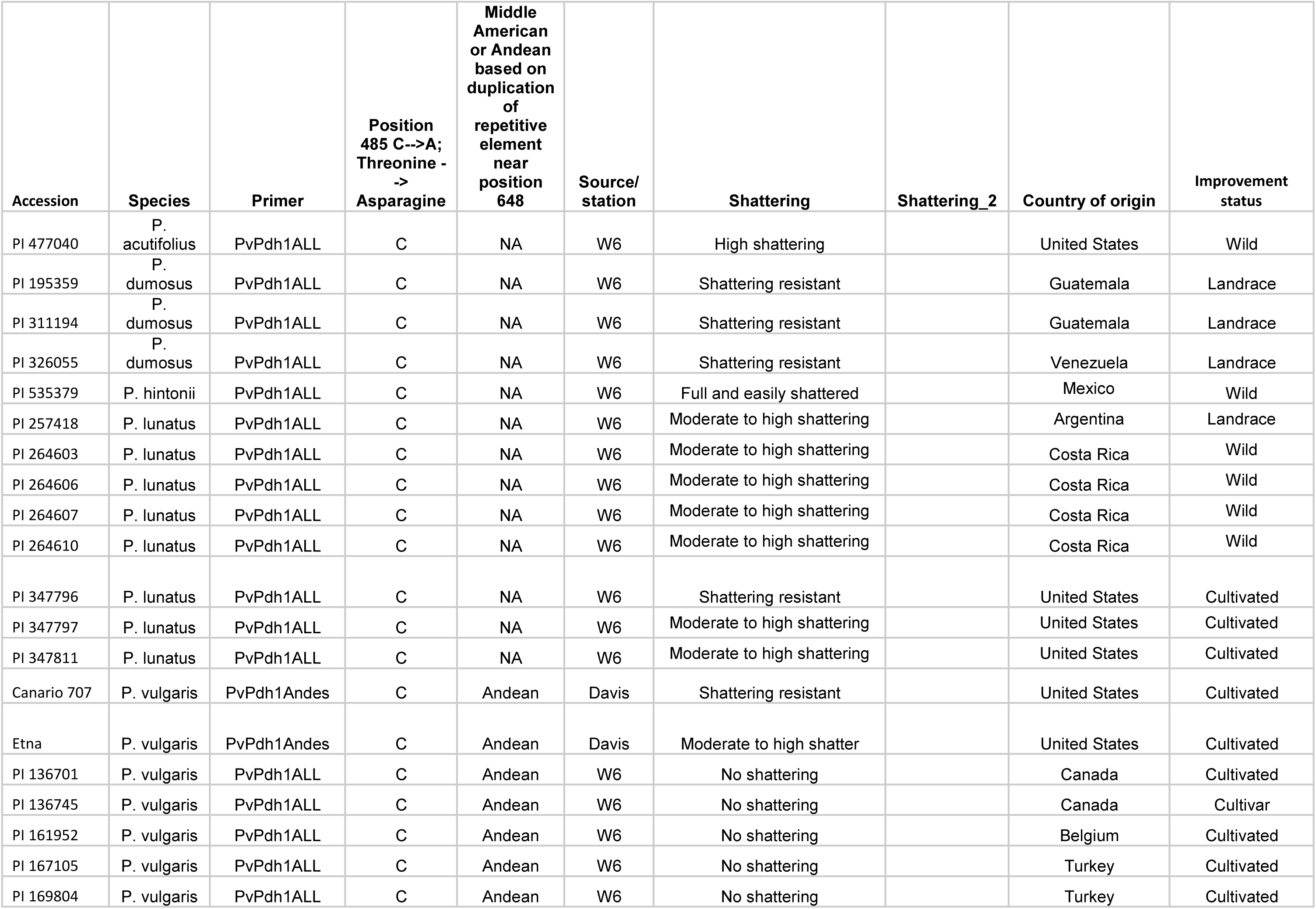
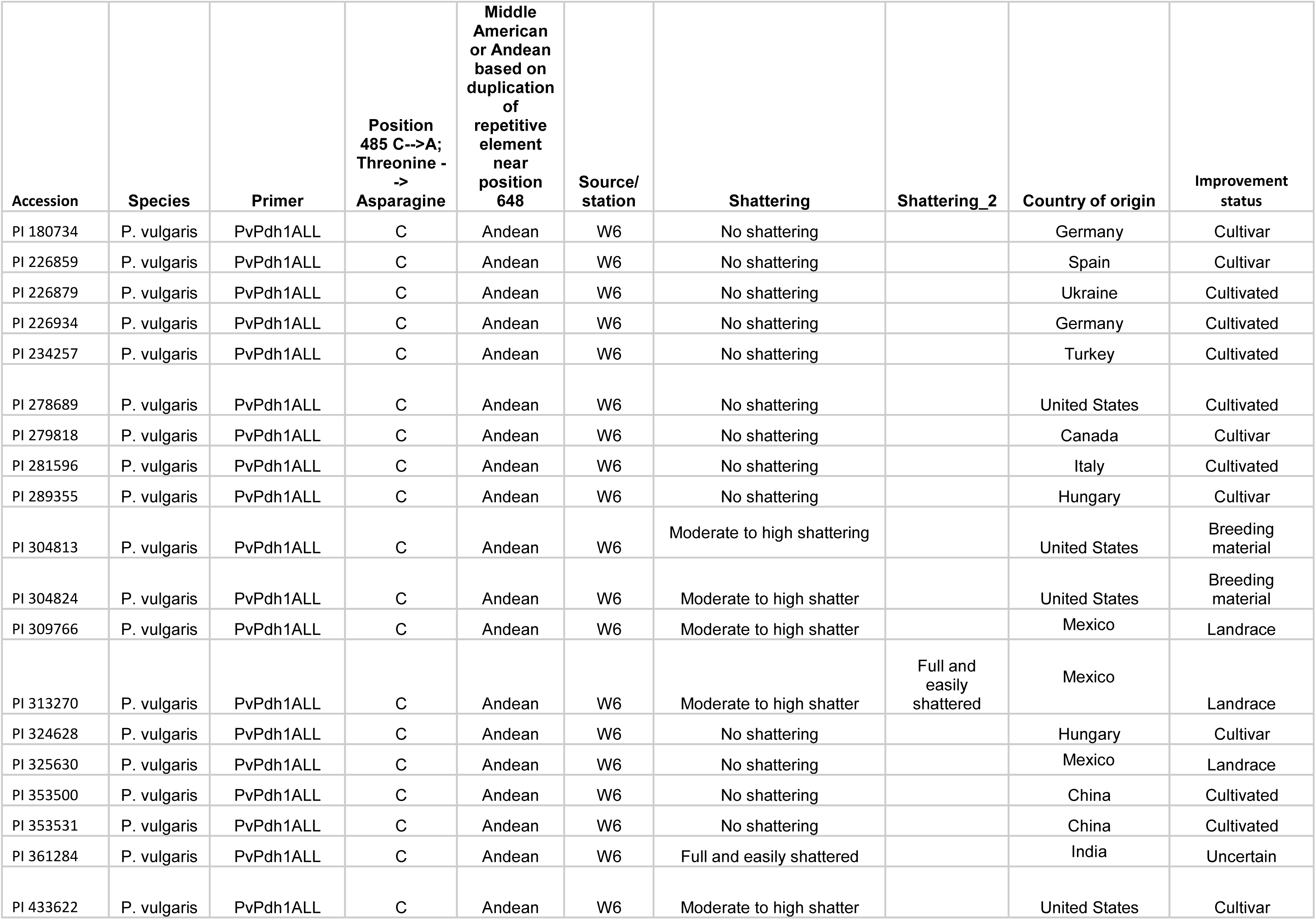
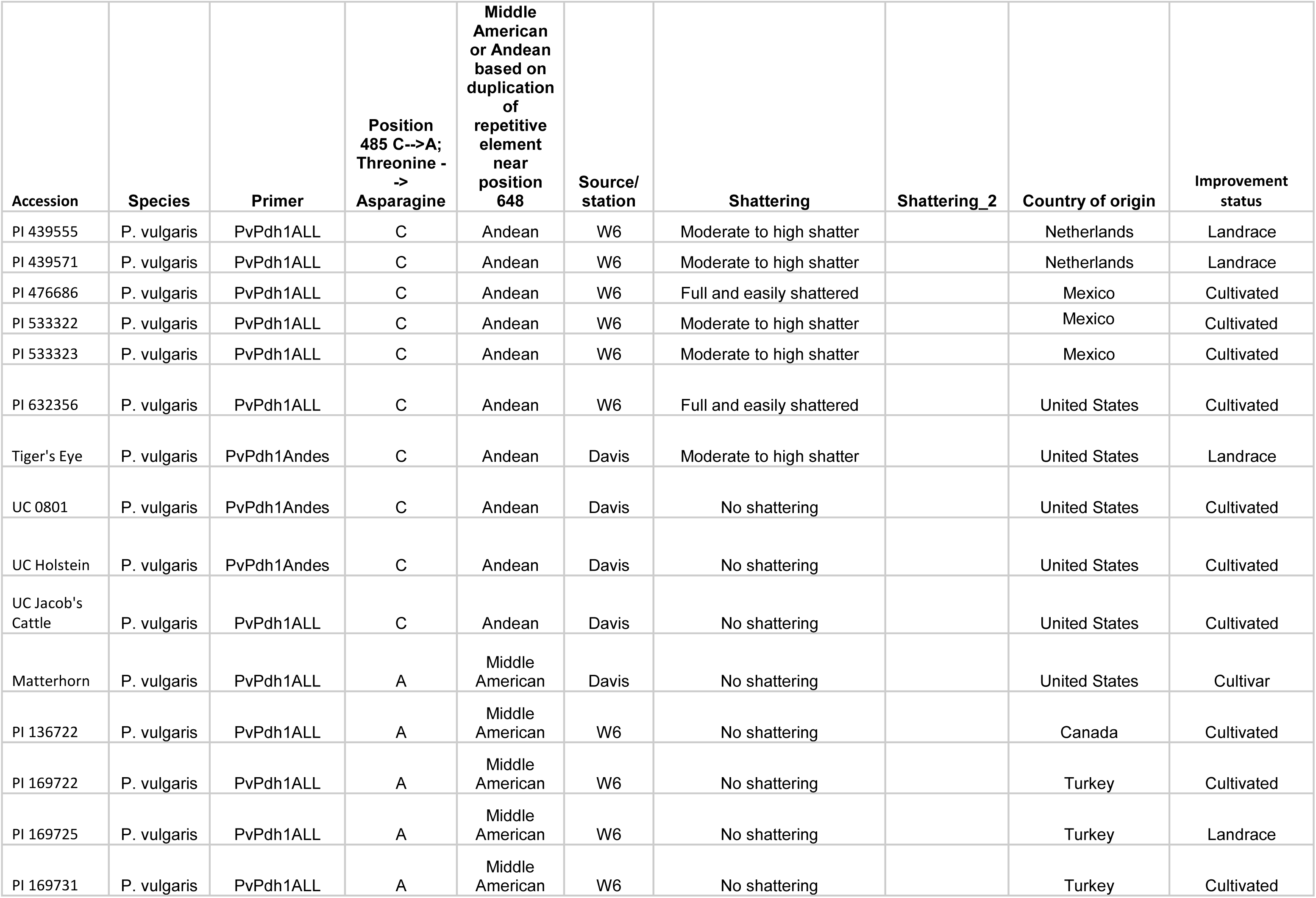
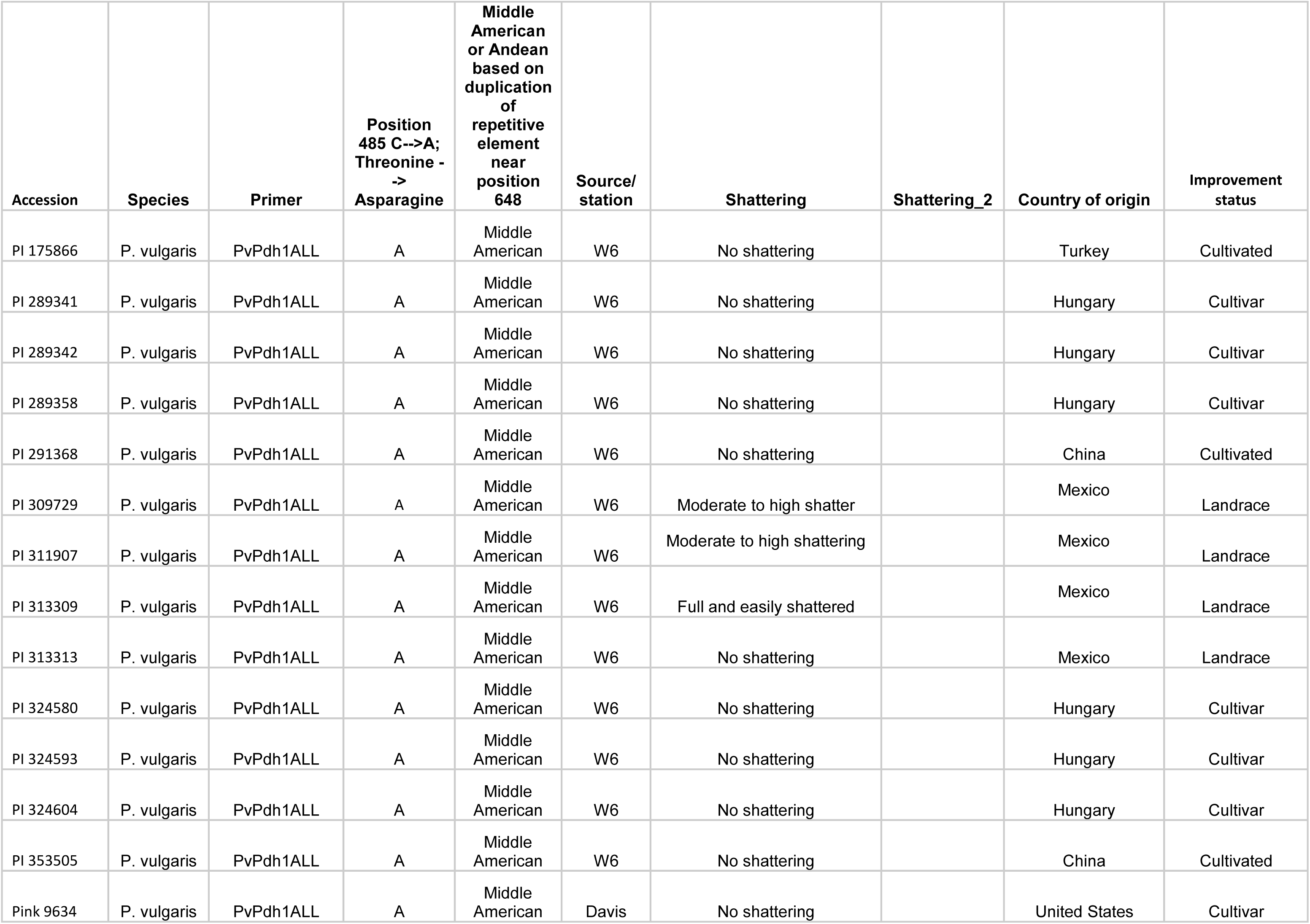
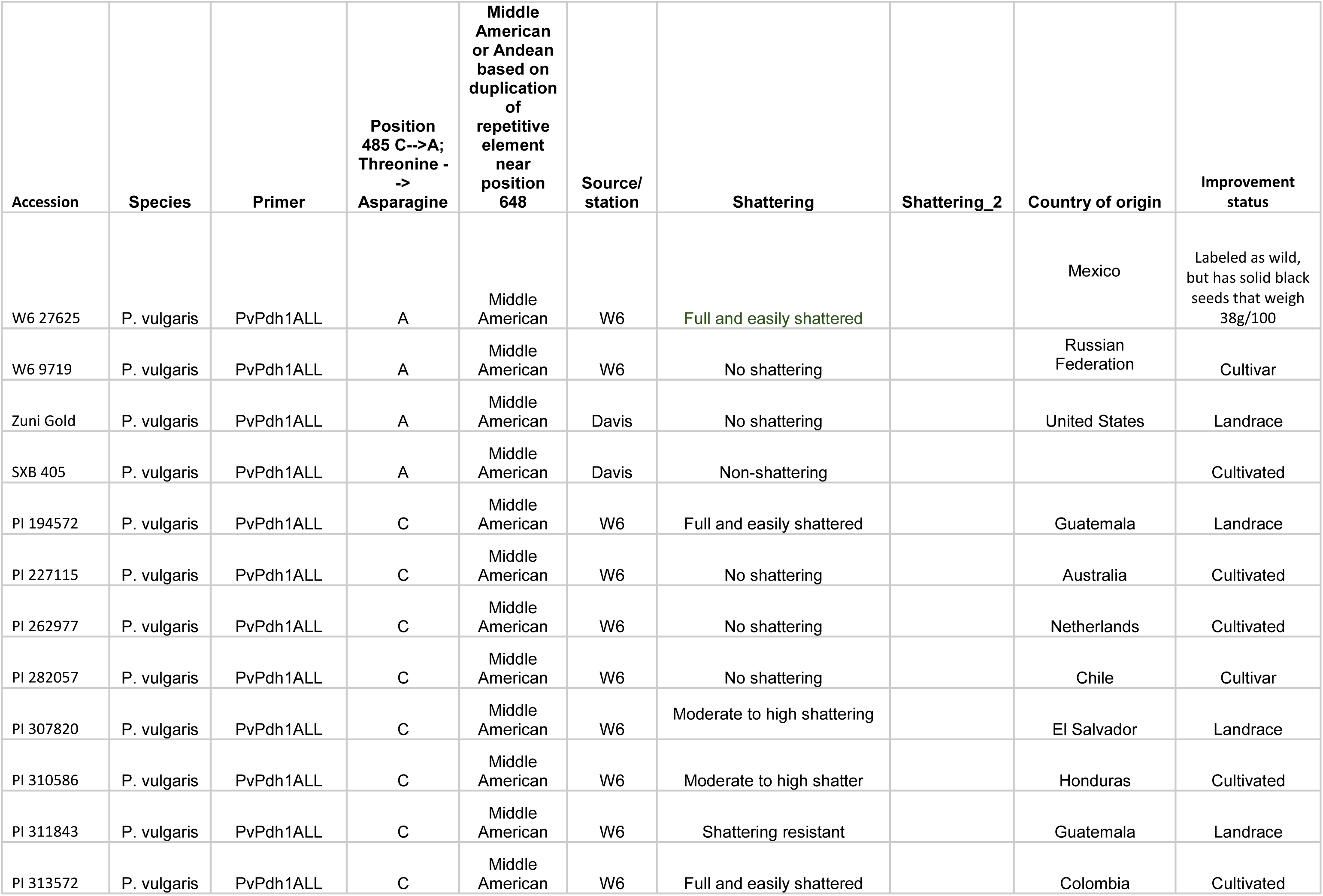
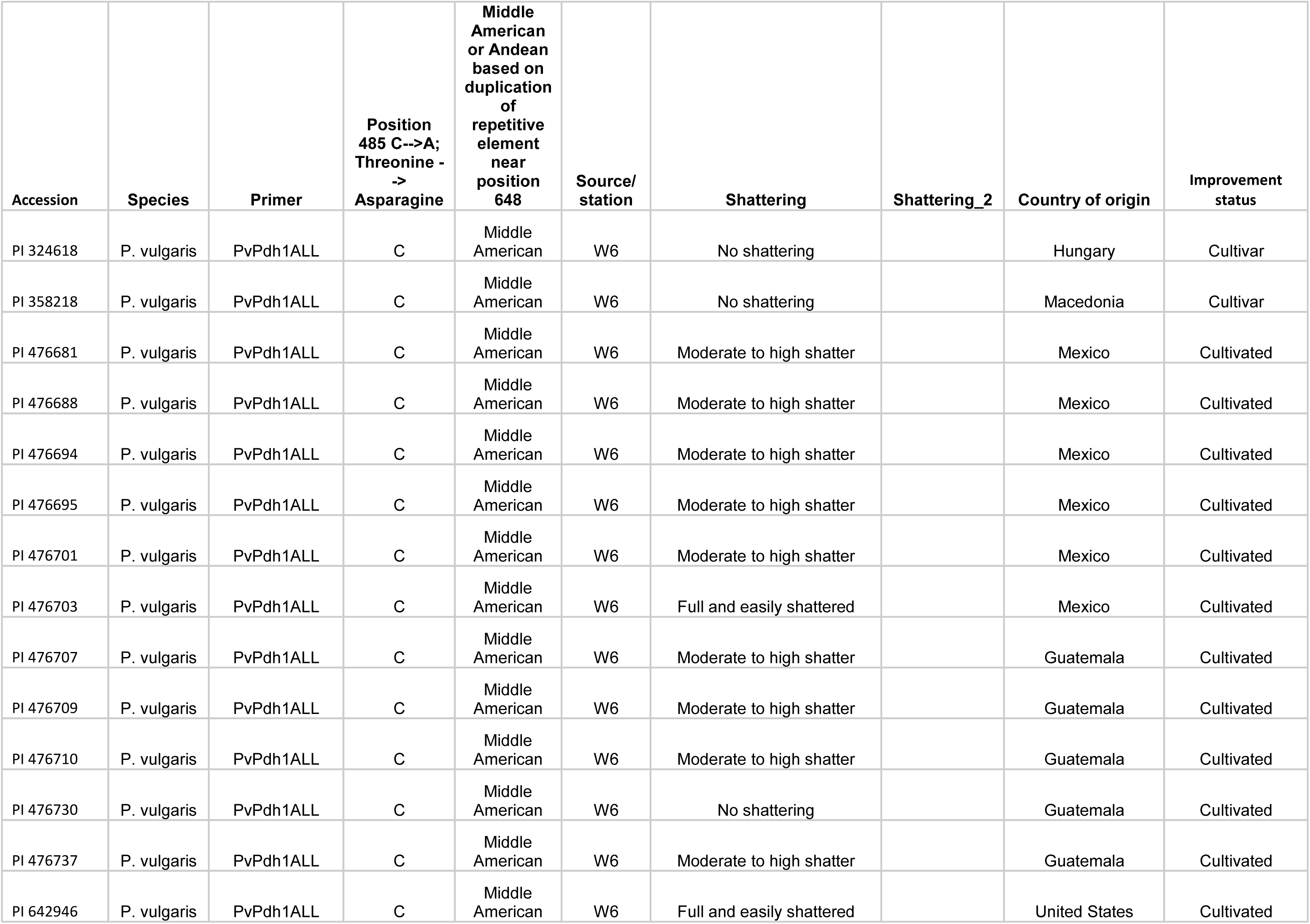
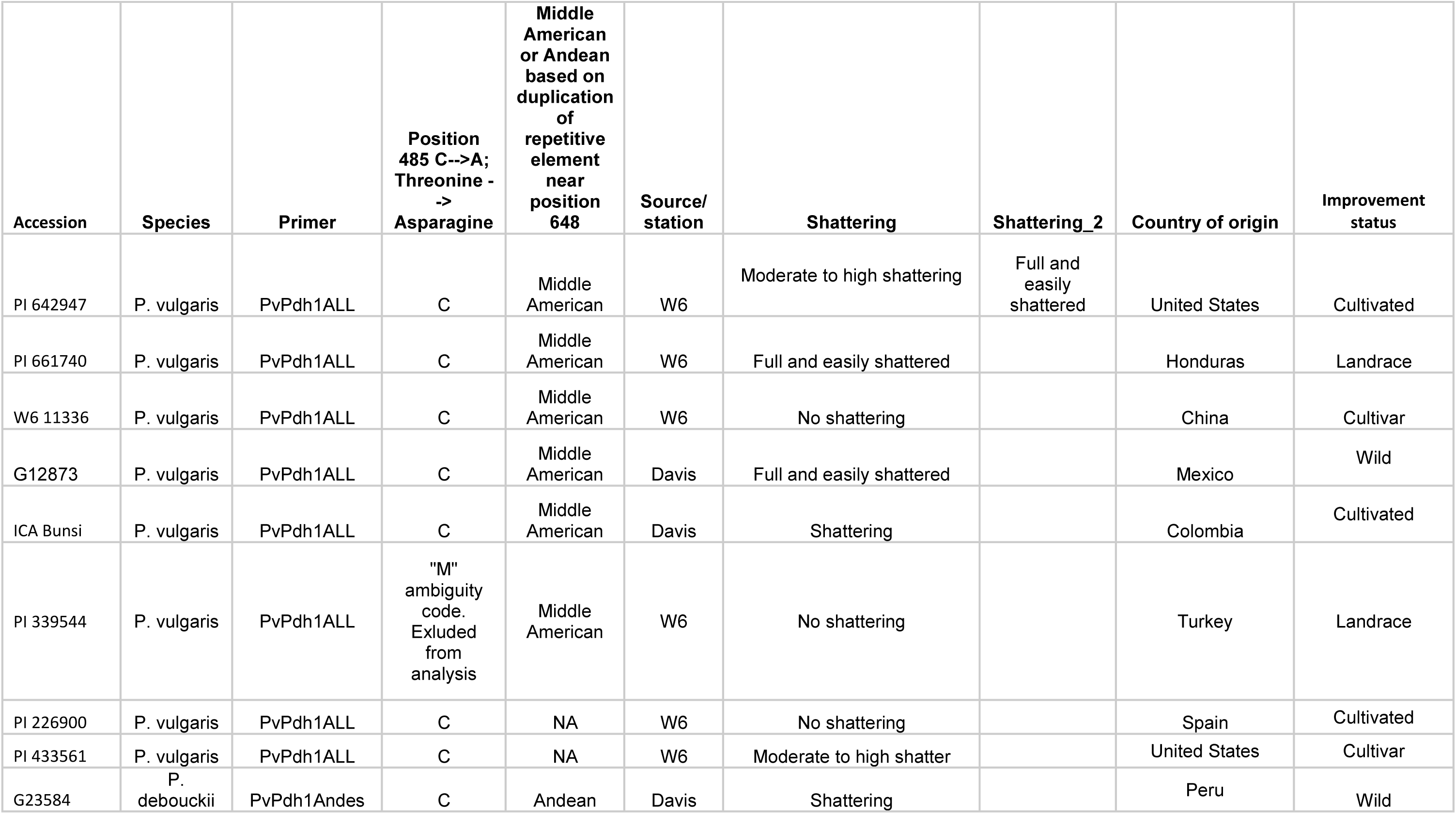
Sequencing of *PvPdh1* in several species of wild and domesticated *Phaseolus*

| Accession | Species | Primer | Position<br>485 C-->A;<br>Threonine -<br>-><br>Asparagine | Middle<br>American<br>or Andean<br>based on<br>duplication<br>of<br>repetitive<br>element<br>near<br>position<br>648 | Source/<br>station | Shattering | Shattering_2 | Country of origin | Improvement<br>status |
| --- | --- | --- | --- | --- | --- | --- | --- | --- | --- |
| PI 477040 | *P.*<br>*acutifolius* | PvPdh1ALL | C | NA | W6 | High shattering |  | United States | Wild |
| PI 195359 | *P.*<br>*dumosus* | PvPdh1ALL | C | NA | W6 | Shattering resistant |  | Guatemala | Landrace |
| PI 311194 | *P.*<br>*dumosus* | PvPdh1ALL | C | NA | W6 | Shattering resistant |  | Guatemala | Landrace |
| PI 326055 | *P.*<br>*dumosus* | PvPdh1ALL | C | NA | W6 | Shattering resistant |  | Venezuela | Landrace |
| PI 535379 | *P. hintonii* | PvPdh1ALL | C | NA | W6 | Full and easily shattered |  | Mexico | Wild |
| PI 257418 | *P. lunatus* | PvPdh1ALL | C | NA | W6 | Moderate to high shattering |  | Argentina | Landrace |
| PI 264603 | *P. lunatus* | PvPdh1ALL | C | NA | W6 | Moderate to high shattering |  | Costa Rica | Wild |
| PI 264606 | *P. lunatus* | PvPdh1ALL | C | NA | W6 | Moderate to high shattering |  | Costa Rica | Wild |
| PI 264607 | *P. lunatus* | PvPdh1ALL | C | NA | W6 | Moderate to high shattering |  | Costa Rica | Wild |
| PI 264610 | *P. lunatus* | PvPdh1ALL | C | NA | W6 | Moderate to high shattering |  | Costa Rica | Wild |
| PI 347796 | *P. lunatus* | PvPdh1ALL | C | NA | W6 | Shattering resistant |  | United States | Cultivated |
| PI 347797 | *P. lunatus* | PvPdh1ALL | C | NA | W6 | Moderate to high shattering |  | United States | Cultivated |
| PI 347811 | *P. lunatus* | PvPdh1ALL | C | NA | W6 | Moderate to high shattering |  | United States | Cultivated |
| Canario 707 | *P. vulgaris* | PvPdh1Andes | C | Andean | Davis | Shattering resistant |  | United States | Cultivated |
| Etna | *P. vulgaris* | PvPdh1Andes | C | Andean | Davis | Moderate to high shatter |  | United States | Cultivated |
| PI 136701 | *P. vulgaris* | PvPdh1ALL | C | Andean | W6 | No shattering |  | Canada | Cultivated |
| PI 136745 | *P. vulgaris* | PvPdh1ALL | C | Andean | W6 | No shattering |  | Canada | Cultivar |
| PI 161952 | *P. vulgaris* | PvPdh1ALL | C | Andean | W6 | No shattering |  | Belgium | Cultivated |
| PI 167105 | *P. vulgaris* | PvPdh1ALL | C | Andean | W6 | No shattering |  | Turkey | Cultivated |
| PI 169804 | *P. vulgaris* | PvPdh1ALL | C | Andean | W6 | No shattering |  | Turkey | Cultivated |
| PI 180734 | P. vulgaris | PvPdh1ALL | C | Andean | W6 | No shattering |  | Germany | Cultivar |
| PI 226859 | P. vulgaris | PvPdh1ALL | C | Andean | W6 | No shattering |  | Spain | Cultivar |
| PI 226879 | P. vulgaris | PvPdh1ALL | C | Andean | W6 | No shattering |  | Ukraine | Cultivated |
| PI 226934 | P. vulgaris | PvPdh1ALL | C | Andean | W6 | No shattering |  | Germany | Cultivated |
| PI 234257 | P. vulgaris | PvPdh1ALL | C | Andean | W6 | No shattering |  | Turkey | Cultivated |
| PI 278689 | P. vulgaris | PvPdh1ALL | C | Andean | W6 | No shattering |  | United States | Cultivated |
| PI 279818 | P. vulgaris | PvPdh1ALL | C | Andean | W6 | No shattering |  | Canada | Cultivar |
| PI 281596 | P. vulgaris | PvPdh1ALL | C | Andean | W6 | No shattering |  | Italy | Cultivated |
| PI 289355 | P. vulgaris | PvPdh1ALL | C | Andean | W6 | No shattering |  | Hungary | Cultivar |
| PI 304813 | P. vulgaris | PvPdh1ALL | C | Andean | W6 | Moderate to high shattering |  | United States | Breeding<br>material |
| PI 304824 | P. vulgaris | PvPdh1ALL | C | Andean | W6 | Moderate to high shatter |  | United States | Breeding<br>material |
| PI 309766 | P. vulgaris | PvPdh1ALL | C | Andean | W6 | Moderate to high shatter |  | Mexico | Landrace |
| PI 313270 | P. vulgaris | PvPdh1ALL | C | Andean | W6 | Moderate to high shatter | Full and<br>easily<br>shattered | Mexico | Landrace |
| PI 324628 | P. vulgaris | PvPdh1ALL | C | Andean | W6 | No shattering |  | Hungary | Cultivar |
| PI 325630 | P. vulgaris | PvPdh1ALL | C | Andean | W6 | No shattering |  | Mexico | Landrace |
| PI 353500 | P. vulgaris | PvPdh1ALL | C | Andean | W6 | No shattering |  | China | Cultivated |
| PI 353531 | P. vulgaris | PvPdh1ALL | C | Andean | W6 | No shattering |  | China | Cultivated |
| PI 361284 | P. vulgaris | PvPdh1ALL | C | Andean | W6 | Full and easily shattered |  | India | Uncertain |
| PI 433622 | P. vulgaris | PvPdh1ALL | C | Andean | W6 | Moderate to high shatter |  | United States | Cultivar |
| PI 439555 | P. vulgaris | PvPdh1ALL | C | Andean | W6 | Moderate to high shatter |  | Netherlands | Landrace |
| PI 439571 | P. vulgaris | PvPdh1ALL | C | Andean | W6 | Moderate to high shatter |  | Netherlands | Landrace |
| PI 476686 | P. vulgaris | PvPdh1ALL | C | Andean | W6 | Full and easily shattered |  | Mexico | Cultivated |
| PI 533322 | P. vulgaris | PvPdh1ALL | C | Andean | W6 | Moderate to high shatter |  | Mexico | Cultivated |
| PI 533323 | P. vulgaris | PvPdh1ALL | C | Andean | W6 | Moderate to high shatter |  | Mexico | Cultivated |
| PI 632356 | P. vulgaris | PvPdh1ALL | C | Andean | W6 | Full and easily shattered |  | United States | Cultivated |
| Tiger's Eye | P. vulgaris | PvPdh1Andes | C | Andean | Davis | Moderate to high shatter |  | United States | Landrace |
| UC 0801 | P. vulgaris | PvPdh1Andes | C | Andean | Davis | No shattering |  | United States | Cultivated |
| UC Holstein | P. vulgaris | PvPdh1Andes | C | Andean | Davis | No shattering |  | United States | Cultivated |
| UC Jacob's<br>Cattle | P. vulgaris | PvPdh1ALL | C | Andean | Davis | No shattering |  | United States | Cultivated |
| Matterhorn | P. vulgaris | PvPdh1ALL | A | Middle<br>American | Davis | No shattering |  | United States | Cultivar |
| PI 136722 | P. vulgaris | PvPdh1ALL | A | Middle<br>American | W6 | No shattering |  | Canada | Cultivated |
| PI 169722 | P. vulgaris | PvPdh1ALL | A | Middle<br>American | W6 | No shattering |  | Turkey | Cultivated |
| PI 169725 | P. vulgaris | PvPdh1ALL | A | Middle<br>American | W6 | No shattering |  | Turkey | Landrace |
| PI 169731 | P. vulgaris | PvPdh1ALL | A | Middle<br>American | W6 | No shattering |  | Turkey | Cultivated |

| Accession | Species | Primer | Position<br>485 C-->A;<br>Threonine -<br>-> Asparagine | Middle<br>American<br>or Andean<br>based on<br>duplication<br>of<br>repetitive<br>element<br>near<br>position<br>648 | Source/<br>station | Shattering | Shattering_2 | Country of origin | Improvement<br>status |
| --- | --- | --- | --- | --- | --- | --- | --- | --- | --- |
| PI 175866 | P. vulgaris | PvPdh1ALL | A | Middle<br>American | W6 | No shattering |  | Turkey | Cultivated |
| PI 289341 | P. vulgaris | PvPdh1ALL | A | Middle<br>American | W6 | No shattering |  | Hungary | Cultivar |
| PI 289342 | P. vulgaris | PvPdh1ALL | A | Middle<br>American | W6 | No shattering |  | Hungary | Cultivar |
| PI 289358 | P. vulgaris | PvPdh1ALL | A | Middle<br>American | W6 | No shattering |  | Hungary | Cultivar |
| PI 291368 | P. vulgaris | PvPdh1ALL | A | Middle<br>American | W6 | No shattering |  | China | Cultivated |
| PI 309729 | P. vulgaris | PvPdh1ALL | A | Middle<br>American | W6 | Moderate to high shatter |  | Mexico | Landrace |
| PI 311907 | P. vulgaris | PvPdh1ALL | A | Middle<br>American | W6 | Moderate to high shattering |  | Mexico | Landrace |
| PI 313309 | P. vulgaris | PvPdh1ALL | A | Middle<br>American | W6 | Full and easily shattered |  | Mexico | Landrace |
| PI 313313 | P. vulgaris | PvPdh1ALL | A | Middle<br>American | W6 | No shattering |  | Mexico | Landrace |
| PI 324580 | P. vulgaris | PvPdh1ALL | A | Middle<br>American | W6 | No shattering |  | Hungary | Cultivar |
| PI 324593 | P. vulgaris | PvPdh1ALL | A | Middle<br>American | W6 | No shattering |  | Hungary | Cultivar |
| PI 324604 | P. vulgaris | PvPdh1ALL | A | Middle<br>American | W6 | No shattering |  | Hungary | Cultivar |
| PI 353505 | P. vulgaris | PvPdh1ALL | A | Middle<br>American | W6 | No shattering |  | China | Cultivated |
| Pink 9634 | P. vulgaris | PvPdh1ALL | A | Middle<br>American | Davis | No shattering |  | United States | Cultivar |
| W6 27625 | P. vulgaris | PvPdh1ALL | A | Middle<br>American | W6 | Full and easily shattered |  | Mexico | Labeled as wild,<br>but has solid black<br>seeds that weigh<br>38g/100 |
| W6 9719 | P. vulgaris | PvPdh1ALL | A | Middle<br>American | W6 | No shattering |  | Russian<br>Federation | Cultivar |
| Zuni Gold | P. vulgaris | PvPdh1ALL | A | Middle<br>American | Davis | No shattering |  | United States | Landrace |
| SXB 405 | P. vulgaris | PvPdh1ALL | A | Middle<br>American | Davis | Non-shattering |  |  | Cultivated |
| PI 194572 | P. vulgaris | PvPdh1ALL | C | Middle<br>American | W6 | Full and easily shattered |  | Guatemala | Landrace |
| PI 227115 | P. vulgaris | PvPdh1ALL | C | Middle<br>American | W6 | No shattering |  | Australia | Cultivated |
| PI 262977 | P. vulgaris | PvPdh1ALL | C | Middle<br>American | W6 | No shattering |  | Netherlands | Cultivated |
| PI 282057 | P. vulgaris | PvPdh1ALL | C | Middle<br>American | W6 | No shattering |  | Chile | Cultivar |
| PI 307820 | P. vulgaris | PvPdh1ALL | C | Middle<br>American | W6 | Moderate to high shattering |  | El Salvador | Landrace |
| PI 310586 | P. vulgaris | PvPdh1ALL | C | Middle<br>American | W6 | Moderate to high shatter |  | Honduras | Cultivated |
| PI 311843 | P. vulgaris | PvPdh1ALL | C | Middle<br>American | W6 | Shattering resistant |  | Guatemala | Landrace |
| PI 313572 | P. vulgaris | PvPdh1ALL | C | Middle<br>American | W6 | Full and easily shattered |  | Colombia | Cultivated |
| PI 324618 | P. vulgaris | PvPdh1ALL | C | Middle<br>American | W6 | No shattering |  | Hungary | Cultivar |
| PI 358218 | P. vulgaris | PvPdh1ALL | C | Middle<br>American | W6 | No shattering |  | Macedonia | Cultivar |
| PI 476681 | P. vulgaris | PvPdh1ALL | C | Middle<br>American | W6 | Moderate to high shatter |  | Mexico | Cultivated |
| PI 476688 | P. vulgaris | PvPdh1ALL | C | Middle<br>American | W6 | Moderate to high shatter |  | Mexico | Cultivated |
| PI 476694 | P. vulgaris | PvPdh1ALL | C | Middle<br>American | W6 | Moderate to high shatter |  | Mexico | Cultivated |
| PI 476695 | P. vulgaris | PvPdh1ALL | C | Middle<br>American | W6 | Moderate to high shatter |  | Mexico | Cultivated |
| PI 476701 | P. vulgaris | PvPdh1ALL | C | Middle<br>American | W6 | Moderate to high shatter |  | Mexico | Cultivated |
| PI 476703 | P. vulgaris | PvPdh1ALL | C | Middle<br>American | W6 | Full and easily shattered |  | Mexico | Cultivated |
| PI 476707 | P. vulgaris | PvPdh1ALL | C | Middle<br>American | W6 | Moderate to high shatter |  | Guatemala | Cultivated |
| PI 476709 | P. vulgaris | PvPdh1ALL | C | Middle<br>American | W6 | Moderate to high shatter |  | Guatemala | Cultivated |
| PI 476710 | P. vulgaris | PvPdh1ALL | C | Middle<br>American | W6 | Moderate to high shatter |  | Guatemala | Cultivated |
| PI 476730 | P. vulgaris | PvPdh1ALL | C | Middle<br>American | W6 | No shattering |  | Guatemala | Cultivated |
| PI 476737 | P. vulgaris | PvPdh1ALL | C | Middle<br>American | W6 | Moderate to high shatter |  | Guatemala | Cultivated |
| PI 642946 | P. vulgaris | PvPdh1ALL | C | Middle<br>American | W6 | Full and easily shattered |  | United States | Cultivated |
| PI 642947 | P. vulgaris | PvPdh1ALL | C | Middle<br>American | W6 | Moderate to high shattering | Full and<br>easily<br>shattered | United States | Cultivated |
| PI 661740 | P. vulgaris | PvPdh1ALL | C | Middle<br>American | W6 | Full and easily shattered |  | Honduras | Landrace |
| W6 11336 | P. vulgaris | PvPdh1ALL | C | Middle<br>American | W6 | No shattering |  | China | Cultivar |
| G12873 | P. vulgaris | PvPdh1ALL | C | Middle<br>American | Davis | Full and easily shattered |  | Mexico | Wild |
| ICA Bunsí | P. vulgaris | PvPdh1ALL | C | Middle<br>American | Davis | Shattering |  | Colombia | Cultivated |
| PI 339544 | P. vulgaris | PvPdh1ALL | "M"<br>ambiguity<br>code.<br>Excluded<br>from<br>analysis | Middle<br>American | W6 | No shattering |  | Turkey | Landrace |
| PI 226900 | P. vulgaris | PvPdh1ALL | C | NA | W6 | No shattering |  | Spain | Cultivated |
| PI 433561 | P. vulgaris | PvPdh1ALL | C | NA | W6 | Moderate to high shatter |  | United States | Cultivar |
| G23584 | P.<br>debouckii | PvPdh1Andes | C | Andean | Davis | Shattering |  | Peru | Wild |

**Table S3.** Synteny near *Pdh1* in *G. max* and *P. vulgaris* - sharing of gene models.

| *Glycine max* |  | *Phaseolus vulgaris* |  |
| --- | --- | --- | --- |
| Gene model | Predicted gene function | Gene model | Predicted gene function |
| Glyma16g25490.1 | Proline-rich extensin-like receptor kinase 4 | Phvul.003G252900.1 | Proline-rich extensin-like receptor kinase 4 |
| Glyma16g25500.1 | Leucine-rich repeat (LRR) family protein | Phvul.003G252700.1 | Leucine-rich repeat (LRR) family protein |
| Glyma16g25530.2 | Plant invertase/pectin methylesterase inhibitor superfamily protein |  |  |
| Glyma16g25540.1 | Major facilitator superfamily protein | Phvul.003G252500.1 | Major facilitator superfamily protein |
| Glyma16g25550.1 | C2H2-like zinc finger protein | Phvul.003G252400.1 | C2H2-like zinc finger protein |
| Glyma16g25560.1 | Protein phosphatase 2C family protein | Phvul.003G252300.1 | Protein phosphatase 2C family protein |
| Glyma16g25570.1 | GRAS family transcription factor | Phvul.003G252200.1 | GRAS family transcription factor |
| **Glyma16g25580.1** | **Disease resistance-responsive (dirigent-like protein) family protein** | **Phvul.003G252100.1** | **Disease resistance-responsive (dirigent-like protein) family protein** |
| Glyma16g25600.1 | G-box binding factor 1 |  |  |
| Glyma16g25600.5 | G-box binding factor 1 | Phvul.003G252000.1 | G-box binding factor 1 |
| Glyma16g25600.4 | G-box binding factor 1 |  |  |
| Glyma16g25600.6 | G-box binding factor 1 |  |  |
| Glyma16g25600.7 | G-box binding factor 1 |  |  |
| Glyma16g25611.1 | Protein kinase superfamily protein | Phvul.003G251900.1 | Protein kinase superfamily protein |
| Glyma16g25620.1 | geranylgeranyl pyrophosphate synthase 1 | Phvul.003G251500.1 | geranylgeranyl pyrophosphate synthase 1 |
| Glyma16g25650.1 | Protein of unknown function (DUF677) |  |  |
| Glyma16g25660.1 |  |  |  |
| Glyma16g25670.1 |  |  |  |
| Glyma16g25680.2 | Protein of unknown function (DUF607) | Phvul.003G251400.1 | Protein of unknown function (DUF607) |
| Glyma16g25690.1 | GNS1/SUR4 membrane protein family | Phvul.003G251300.1 | GNS1/SUR4 membrane protein family |
| Glyma16g25700.1 | PQ-loop repeat family protein / transmembrane family protein | Phvul.003G251100.1 | PQ-loop repeat family protein / transmembrane family protein |
| Glyma16g25710.2 | Pectin lyase-like superfamily protein |  |  |
| Glyma16g25720.1 | calmodulin-like 41 | Phvul.003G251000.1 | calmodulin-like 41 |
| Glyma16g25740.2 | vacuolar ATP synthase subunit H family protein | Phvul.003G250900.1 | vacuolar ATP synthase subunit H family protein |

**Table S4.** PD after desiccation, by market class, gene pool, and ecogeographic race

| Market class | Gene pool | Race | Mean PD (%) | Median PD (%) | Standard deviation (%) | n |
| --- | --- | --- | --- | --- | --- | --- |
| Cranberry | Andean | Nueva Granada | 41.43 | 46.29 | 29.86 | 24 |
| Kidney | Andean | Nueva Granada | 21.09 | 13.89 | 18.32 | 43 |
| Purple speck/mottled | Andean | Nueva Granada | 3.11 | 0 | 5.84 | 17 |
| Great northern | Middle American | Durango | 0.94 | 0 | 2.12 | 31 |
| Pink | Middle American | Durango | 2.48 | 0 | 6.37 | 23 |
| Pinto | Middle American | Durango | 0.74 | 0 | 2.38 | 93 |
| Black | Middle American | Mesoamerica | 17.63 | 19 | 13.22 | 43 |
| Navy/small white | Middle American | Mesoamerica | 15.2 | 8.5 | 16.62 | 46 |
| Red/small red | Middle American | Variable | 9.59 | 4 | 14.7 | 29 |

Author contributions: T.A.P. prepared the manuscript and conductedlaboratory phenotyping, QTL mapping, GWAS, microscopy, and sequencing. J.C.B.M.T. genotypedthe IxS population, gathered field phenotypes, co-conducted QTL mapping, and provided guidancefor other procedures. A.P. assisted with field and greenhouse trials. J.J. led the sectioning and microscopy studies. P.G.conceived the initial project and provided guidance. All authors edited the manuscript

